# Recombinant production of spider silk protein in Physcomitrella photobioreactors

**DOI:** 10.1101/2024.11.27.625602

**Authors:** Maryam Ramezaniaghdam, Lennard L. Bohlender, Juliana Parsons, Sebastian N. W. Hoernstein, Eva L. Decker, Ralf Reski

## Abstract

Spider dragline silk stands out as a remarkable biomaterial, representing one of nature’s toughest fibres. Its strength rivals that of many synthetic fibres used commercially, rendering it applicable across various industrial and medical domains. However, its widespread utilisation requires cost-effective mass production. Biotechnology presents a promising avenue for achieving this goal, particularly through the production of recombinant dragline silk proteins in transgenic plant systems. This study aimed to assess the feasibility of producing one key protein component of dragline silk, MaSp1, from the western black widow spider, *Latrodectus hesperus*, (LhMaSp1) in the moss Physcomitrella (*Physcomitrium patens*). Here, we present the successful recombinant production of spider silk protein containing both the N- and C-terminal domains (NTD and CTD) of LhMaSp1 protein in moss cells. NTD and CTD are necessary domains for protein assembly in spider silk. The production of recombinant LhMaSp1 protein in Physcomitrella was performed in shake flasks and in five-litre photobioreactors and the correct synthesis of LhMaSp1 was proven *via* mass spectrometry.

**Key message:** We report the first successful plant-produced recombinant spider silk key protein component containing both the N- and the C-terminal domain.

## Introduction

Web-spinning spiders are arachnids that possess the unique ability to produce various types of silk, each tailored to serve a specific purpose. This versatile silk production allows them to efficiently trap prey, construct protective egg cases, evade predators, and secure themselves to surfaces (Vollrath and Knight 2001). Their unique combination of high tensile strength and extensibility results in an extraordinary toughness that inspires research and innovation (Gosline et al. 1999; Qin et al. 2024). It offers a wide range of potential applications. In industry, its durability and elasticity open up possibilities for creating high-performance materials for textiles, composites (Kucharczyk et al. 2019), and biodegradable plastics (Humenik et al. 2011). Spider silk proteins are also biocompatible, supporting cell growth and integration. Because of these properties, spider silks are regarded as a promising material for medical applications such as wound healing (Altman et al. 2003; Öksüz et al. 2021), degradable biosensors for biomonitoring of analytes in the body (Xu et al. 2019), tissue engineering (Salehi et al. 2020), such as artificial blood vessels (Dastagir et al. 2020), nerve regeneration (Kornfeld et al. 2021; Millesi et al. 2021), and scaffolds creation (Gellynck et al. 2008). They do not provoke immune responses, making them safe for use in humans without causing adverse reactions (Gellynck et al. 2008). These unique properties have sparked considerable interest in the potential applications of spider dragline silk across various fields.

Among the different kinds of silk, dragline silk stands out for its combination of strength and elasticity. This silk’s mechanical properties enable it to absorb and dissipate substantial energy, making it exceptionally resilient (Gosline et al. 1999; Malay et al. 2017). The mechanical properties of dragline silk arise from the structural organisation of proteins called spidroins. The primary components of dragline silk are major ampullate spidroins (MaSp) 1 and 2. These large proteins, which range in size from 200 kDa to 350 kDa, feature a distinctive arrangement: a central repetitive region flanked by non-repetitive and evolutionarily conserved N-terminal and C-terminal domains (NTD and CTD). The sequence characteristics of dragline repetitive regions differ among spider species (Ramezaniaghdam et al. 2022). Some spidroins have short, simple repeat units, while others consist of longer, more complex repeats (Hayashi et al. 1999; Gatesy et al. 2001; Römer and Scheibel 2008). Both the CTD and NTD play crucial roles in initiating fibre formation. The NTD, in particular, can enhance spidroin solubility and regulate fibre formation through pH-dependent dimerization (Askarieh et al. 2010; Rissing et al. 2010; Hagn et al. 2011), while the CTD can trigger the transition of the repetitive region into β-sheet conformation (De Oliveira et al. 2024).

Spiders are notoriously difficult to farm for their silk due to their predatory and cannibalistic nature, which makes large-scale production impractical. To overcome this, researchers have turned to transgenic technologies to develop biomimetic silks (Bini et al. 2006). This approach has involved various host organisms. Each of these systems offers insights and incremental progress toward the goal of producing spider silk at a scale and a quality suitable for industrial and medical applications, but each also highlights the ongoing challenges (Whittall et al. 2021; Ramezaniaghdam et al. 2022). Spider silks in their native forms contain intrinsically disordered regions and repetitive sequences. When such native sequences are recombinantly expressed in bacteria, they are prone to premature aggregation in inclusion bodies (Rinas et al. 2017). Bacterial systems also encounter the challenge of producing spidroins with multiple molecular weights (Xia et al. 2010; Bowen et al. 2018). Baby hamster kidney (BHK) cells and bovine mammary epithelial alveolar cells (MAC-T) have also been used as hosts for silk protein production (Lazaris et al. 2002). In another approach spider silk proteins were successfully produced in goat milk (Copeland et al. 2015). While mammalian cells and transgenic animals offer a more complex and potentially more suitable environment for protein folding and post-translational modifications, the yield of spider silk proteins in these systems has been low, limiting their practicality for large-scale production (Lazaris et al. 2002; Copeland et al. 2015). Insects have been explored as another potential production system due to the relatively close evolutionary distance between spiders and insects (Huemmerich et al. 2004; Anton et al. 2017). Although they could produce spider silk filaments, the process is time-consuming, which poses a significant barrier to efficient production. Some challenges are identified with using the yeast *Pichia pastoris* (new name *Komagataella phaffii*) as a production system for spider silk protein, such as poor expression and proteolysis (Werten et al. 2019). Plant systems such as tobacco (Menassa et al. 2004; Hauptmann et al. 2013; Weichert et al. 2014) and Arabidopsis (Barr et al. 2004; Yang et al. 2005) have shown successful production of spider silk proteins, albeit at low yields. So far, proteins produced in plant systems lack the crucial terminal domains, which are essential for the assembly, mechanical properties, and functionality of the silk.

The objective of this work was to evaluate the potential of the model moss Physcomitrella (*Physcomitrium patens*; Lueth and Reski 2023) as a production platform for recombinant spider dragline silk proteins with N- and C-terminal domains. Physcomitrella is an established host system for the safe and efficient production of complex recombinant proteins (Decker and Reski 2020). Physcomitrella is exquisitely amenable to precise genome engineering (Wiedemann et al. 2018; Reski et al. 2018) and able to produce difficult-to-express proteins such as human erythropoietin (Parsons et al. 2012) and human factor H (FH) with 150 kDa (Michelfelder et al. 2017). FH has 20 repetitive protein domains, each of which consists of around 60 amino acids. It is a single-chain molecule linked by 40 intramolecular disulfide bridges (Büttner-Mainik et al. 2011). In Physcomitrella, the secretion of the produced proteins to the surrounding medium is possible by using secretory signals (Decker and Reski 2007), facilitating the subsequent recombinant protein purification. In this research, we began by evaluating spider silk protein production transiently and then progressed to stable production in shake flasks, followed by cultivation on bioreactors. We utilized mass spectrometry to confirm the sequence of the produced proteins.

## Results and discussion

### Transient expression of NTD-LhMaSp1-12Rep-CTD-citrine

The major ampullate silk from *Latrodectus hesperus*, commonly known as the western black widow spider, is known for its strength and great extensibility (Lawrence et al. 2004), making the recombinant production of the Major ampullate silk protein 1 (LhMasp1) highly interesting. As protocols for the fast evaluation of recombinant protein production in Physcomitrella protoplasts are well established (Baur et al. 2005; Schaaf et al. 2005), we assessed the expression of a highly repetitive NTD-LhMaSp1-12Rep-CTD protein in this transient expression system.

The coding sequences (CDSs) of 12 repetitive poly-alanine blocks of the *L. hesperus* silk protein core regions, the full sequence of the N-terminal domain and the C-terminal domain were optimised (Supplementary Fig S1) for Physcomitrella expression according to Top et al. (2021) in order to prevent potential hetero-splicing events and to adapt to the Physcomitrella codon usage. This construct was synthesized in the pUC-GW-Amp plasmid (Genewiz, Leipzig, Germany). The CDS of NTD-LhMaSp1-12Rep-CTD was cloned into an expression vector containing PpActin5, the CDS of citrine, and the nos terminator sequence via *Xho*Ⅰ and *Kpn*Ⅰ restriction sites. Gibson assembly (Gibson et al. 2009) was used to include the Physcomitrella aspartic protease signal peptide (APsp; Schaaf et al. 2004). The assembled vector (Supplemental Fig. S2A) was verified by sequencing.

Protoplasts were monitored 2, 4, 7, and 9 days after transfection *via* fluorescence microscopy. The fluorescence microscopy images reveal the successful production of NTD-LhMaSp1-12Rep-CTD-citrine in Physcomitrella protoplasts (Fig. 1). The citrine signal was observed from day 2, and the signal was still present until day 9. In this construct, we employed the aspartic protease signal peptide (APsp), previously demonstrated to effectively direct GFP to the secretory pathway in Physcomitrella (Schaaf et al. 2004). The presence of ramified network-like fluorescence signals, alongside fluorescence accumulation within the nuclear envelope, serves as evidence for protein targeting to the endoplasmic reticulum (ER) (Schaaf et al. 2004). However, structures reminiscent of an ER network were not observed in cells producing NTD-LhMaSp1-12Rep-CTD-citrine. In contrast, we observed accumulation of protein bodies (PBs) of different sizes (Fig. 1). PB-formation is also described to occur in leaves of *Nicotiana benthamiana* when recombinant proteins are transiently produced at high quantities in the ER, or when recombinant proteins are fused to tags such as Zera, elastin-like polypeptides, and hydrophobins (Conley et al. 2009; Gutiérrez et al. 2013; Saberianfar et al. 2015; Saberianfar et al. 2016). These tags include hydrophobic areas which self-assemble into aggregates of fusion tags and their fusion partners (Conley et al. 2009).

**Fig. 1.**
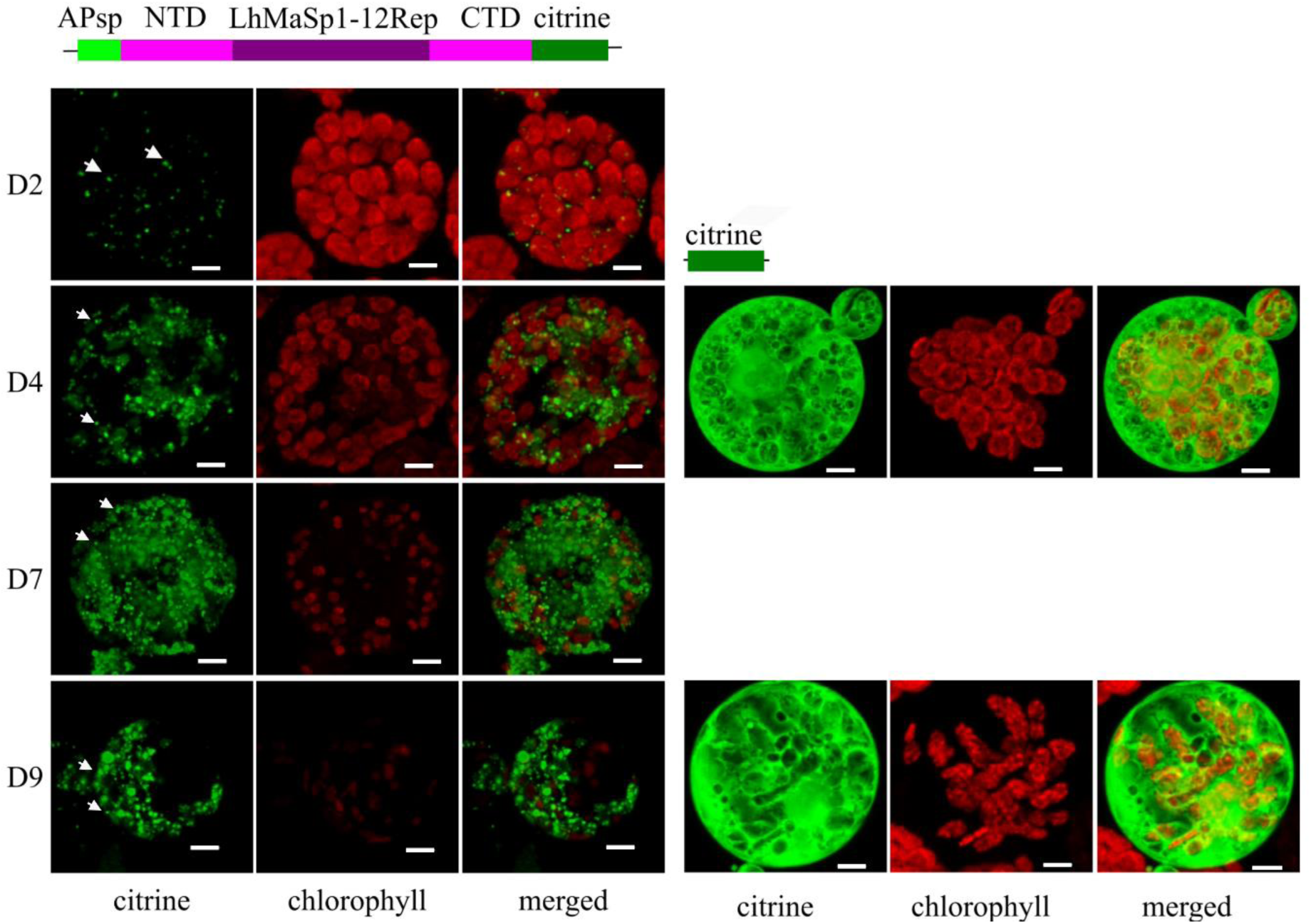
Confocal microscopy images showing the production of NTD-LhMaSp1-12Rep-CTD-citrine fusion protein in Physcomitrella protoplasts. Schematic representation of the fusion protein is at the top. The images were recorded 2 (D2), 4 (D4), 7 (D7), and 9 days (D9) after protoplast transfection and show the successful production of NTD-LhMaSp1-12Rep-CTD-citrine in Physcomitrella. Protein body signals are observed from NTD-LhMaSp1-12Rep-CTD fused to citrine (white arrows). All images are 3D-rendered Z-stack images. NTD: N-terminal domain. CTD: C-terminal domain. Bars = 5 µm.

The secretion of a protein to the extracellular space can facilitate downstream processing (Schaaf et al. 2004) in future experiments. To confirm the presence of NTD-LhMaSp1-12Rep-CTD-citrine in the secretory pathway, a construct encoding mCerulean with an ER retention signal (KDEL) was used to provide an ER marker for co-localisation studies. Physcomitrella protoplasts were co-transfected with both plasmids to investigate potential co-localisation of the respective fluorescent proteins. The 3D rendered Z-stack and single-slice images of protoplasts transiently transformed to produce mCerulean-KDEL and NTD-LhMaSp1-12Rep-CTD-citrine showed the ramified network and the localisation to the nuclear envelope (Fig. 2, Supplementary Fig. S3). The mCerulean marker also showed PBs, albeit they appeared larger than those observed with LhMaSp1. This phenomenon aligns with previous findings in *Nicotiana benthamiana* leaves indicating that secretory pathway proteins can be sequestered into PBs (Saberianfar et al. 2016). Also, protein overexpression may cause artefacts in fluorescence microscopy, including ectopic subcellular localisations, incorrect formation of protein complexes and others (Ratz et al. 2015). The signals emitted by citrine and mCerulean exhibit almost no overlap, albeit demonstrating considerable coverage across similar areas (Fig. 2). A high density of mCerulean-KDEL and NTD-LhMaSp1-12Rep-CTD-citrine signals was observed in an area surrounding the nucleus, which appears to correspond to the rough ER. One possible explanation for the lack of signal overlap could be the formation of PBs at distinct locations within the ER rather than a uniform diffuse signal. Moreover, mCerulean-KDEL is retained in the ER, but NTD-LhMaSp1-12Rep-CTD-citrine can be distributed along the secretory pathway and therefore found in the ER as well as in the Golgi apparatus.

**Fig. 2.**
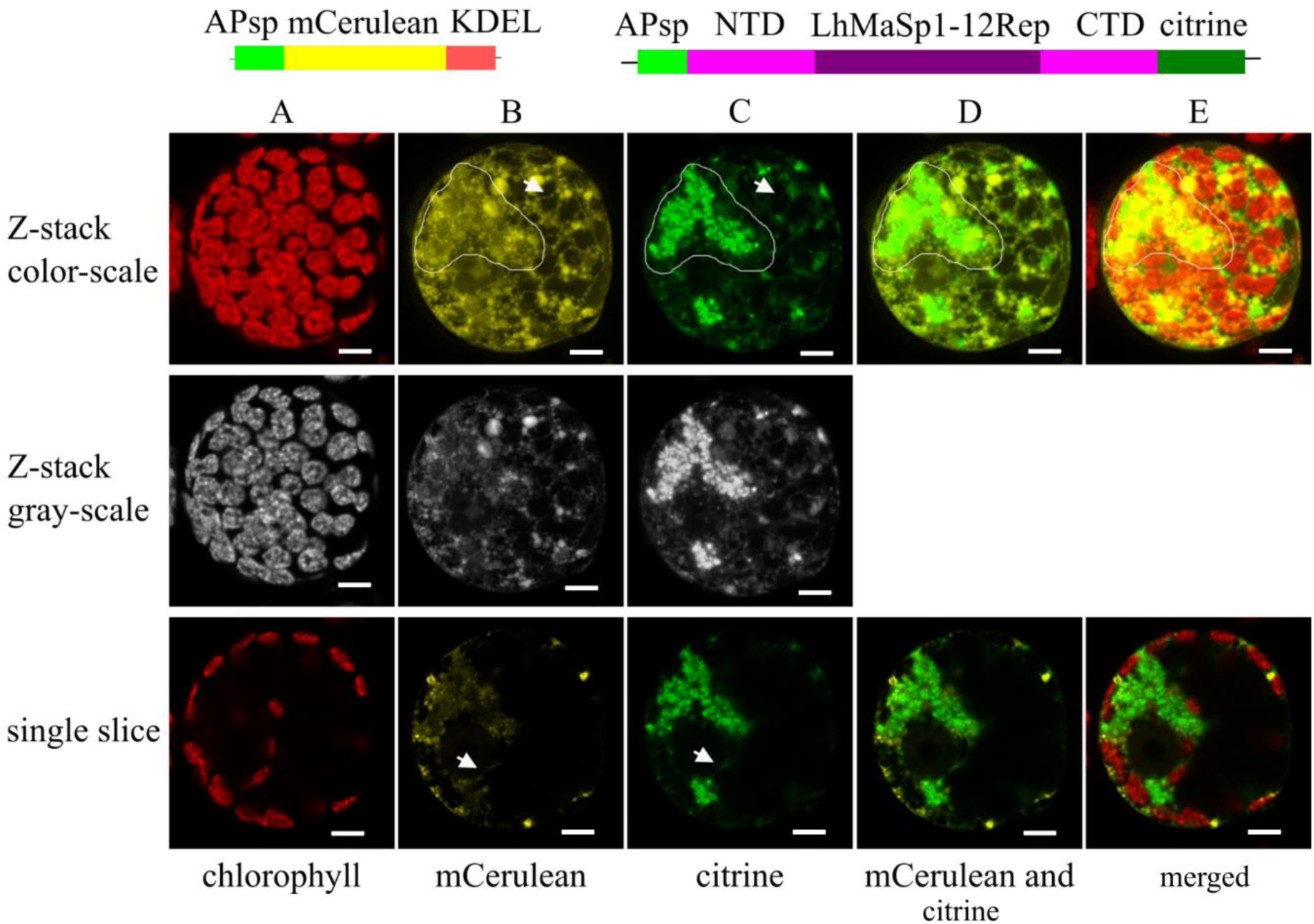
Confocal microscopy images showing the localisation of NTD-LhMaSp1-12Rep-CTD fused to citrine in the secretory pathway of Physcomitrella protoplasts. Schematic representation of the fusion proteins is on top. The mCerulean-KDEL construct was used as a positive control for ER localisation. **A** Chlorophyll autofluorescence. **B, C** mCerulean and citrine channels, respectively. The arrows in Z-stack images point to the fluorescence signal of the ramified network and arrows in single slices point to the fluorescence signal of the nuclear envelope. A dense concentration of mCerulean-KDEL and NTD-LhMaSp1-12Rep-CTD-citrine signals is present in a region encircling the nucleus, suggesting its association with the rough ER. Contour drawings are provided to highlight this area. **D** Citrine and mCerulean signals do not overlap, but demonstrate considerable coverage across similar areas (contour drawings). **E** Merged channels. The upper images display 3D-rendered Z-stack images, while the lower ones depict individual slices of single cells. Grey-scale channels are provided for better visibility. The images were recorded 5 days after protoplast transfection. Bars = 5 µm.

It has been reported that spider silk proteins undergo post-translational modifications (PTMs), including glycosylation (Tillinghast et al. 1992; Sponner et al. 2007; Choresh et al. 2009; Stellwagen and Renberg 2019). However, specific details regarding the type of glycosylation in these proteins remain limited. To date, research on the mechanical characteristics of silk fibres has been conducted with no consideration of the presence of PTMs in the spidroin sequences, although it might influence the mechanical properties of spider silk (Dos Santos-Pinto et al. 2018). Physcomitrella is capable of performing post-translational modifications such as *N*- and *O*-glycosylation (Koprivova et al. 2003; Decker et al. 2014; Bohlender et al. 2022; Stenitzer et al. 2022; Rempfer et al. 2024). The capability of Physcomitrella in transferring proteins to the secretory pathways *via* tailored signal peptides (Decker et al. 2014; Hoernstein et al. 2024) might be helpful in future to produce glycosylated spider silk proteins.

### Stable production of recombinant NTD-LhMaSp1-8Rep-CTD-8Htag

For the stable production of *Latrodectus hesperus* MaSp1 protein in Physcomitrella, the CDSs of eight repetitive poly-alanine blocks of the *L. hesperus* silk protein core region (8Rep), the full sequence of the N-terminal domain (NTD), and the C-terminal domain (CTD) were *in silico* optimised (Supplementary Fig. S4) for expression in Physcomitrella according to Top et al. (2021). The complete construct includes the Physcomitrella actin5 promoter (Weise et al. 2006; Mueller et al. 2014; Niederau et al. 2024), NTD, 8Rep CTD, 8xHis-tag, a nos terminator, and an nptII (neomycin phosphotransferase) antibiotic-resistance expression cassette was synthesized by Genewiz in the pUC-GW-amp plasmid. To have APsp, the construct APsp-NTD-LhMaSp1-12Rep-CTD-8Htag was digested with *Nhe*I and *Eco*RV. The digested product was ligated to the NTD-LhMaSp1-8Rep-CTD**-**8Htag digested with the same enzymes. Assembled vectors (Supplementary Fig. S5A) were verified by sequencing. The schematic representation of the expression cassette for the stable production of NTD-LhMaSp1-8Rep-CTD-8Htag is depicted in Fig. 3A.

**Fig. 3.**
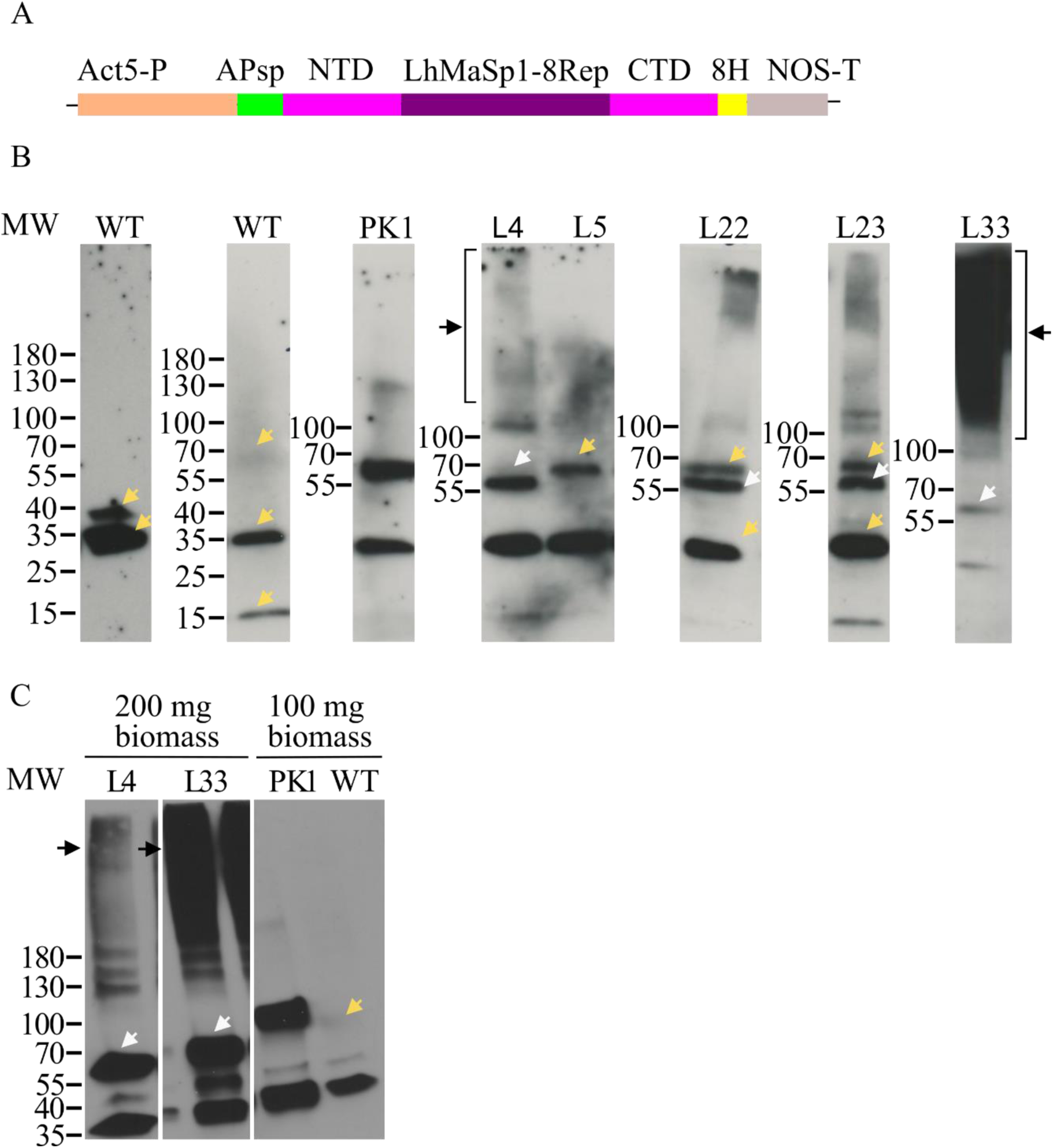
Western blot analysis showing successful production of recombinant NTD-LhMaSp1-8Rep-CTD-8Htag protein in stable Physcomitrella lines. **A** Schematic representation of the expression cassette NTD-LhMaSp1-8Rep-CTD**-**8Htag. The plasmid contains the CDS of aspartic protease signal peptide (APsp), eight codon-optimized repetitive poly-alanine blocks of the *L. hesperus* MaSp1 protein core region, the full sequence of N- and C-terminal domains (NTD and CTD), 8x Histidine-tag (H8), under the control of the Physcomitrella actin5 promoter and the NOS terminator. **B** Immunodetection of NTD-LhMaSp1-8Rep-CTD**-**8Htag was performed *via* Western blot using an anti His-tag antibody (18184, Thermo Fischer Scientific). Some of the best producing lines are presented here. A signal slightly above 55 kDa was detected (white arrows). Almost all transgenic lines show high molecular weight signals higher than 180 kDa (black arrows). **C**. Western blot of the two best NTD-LhMaSp1-8Rep-CTD**-**8Htag producing lines L4 and L33 with the elution fraction from the His SpinTrap column show single bands at ∼56 kDa (white arrows) and high molecular weight signals (black arrows). Unspecific signals also present in the WT are marked with yellow arrows. WT: wild type, L: line. PK1: positive control; transgenic moss line producing the 58 kDa His-tagged protein MFHR1. Gradient 4-15% SDS gels were used. Western blot under reducing conditions. MW: PageRuler Prestained Protein Ladder (Thermo Fisher Scientific).

After transformation with the NTD-LhMaSp1-8Rep-CTD-8Htag construct, the regeneration of 150 transgenic plants was achieved. Fifty transgenic plants were selected for screening with Western blot using an anti His-tag antibody (18184, Thermo Fischer Scientific), and 23 plants with a positive signal were identified (Supplementary Fig. S6). The calculated molecular mass of NTD-LhMaSp1-8Rep-CTD-8Htag without the signal peptide is 52 kDa. The Western blot analysis of some of the best producing lines is presented in Fig. 3. Signals slightly above 55 kDa were observed (Fig. 3B). In addition, almost all transgenic lines showed high molecular weight signals (smear) above 180 kDa (Fig. 3B).

The absence of signals with smaller molecular weight than the expected one in the Western blot suggests that codon optimisation effectively prevented potential heterosplicing events and that NTD-LhMaSp1-8Rep-CTD-8Htag is not degraded in our production conditions. The absence of smaller products indicates the suitability of our expression system, which does not lead to degradation of recombinant MaSp1, typical of some other expression systems such as *Escherichia coli*, *Nicotiana benthamiana*, and *Pichia pastoris* (Fahnestock and Irwin 1997; Menassa et al. 2004; Xia et al. 2010; Bowen et al. 2018).

To assess the purification potential of NTD-LhMaSp1-8Rep-CTD**-**8Htag using the His tag, we used His SpinTrap columns. From 23 positive plants in the first Western blot-based screening, we selected the two lines with the strongest signals: L4, a line which produces mainly a product of slightly above 55 kDa corresponding to the expected molecular weight, and L33, notable for its high molecular weight signals (Fig. 3). Both lines grow similar to the WT on solid medium. For further analysis, the two lines were grown as suspension cultures. Protein extracts from both lines were successfully enriched for NTD-LhMaSp1-8Rep-CTD**-**8Htag using His SpinTrap columns (Fig. 3C, Supplementary Fig. S7), which had no impact of the respective recombinant MaSP1 migration behaviours in SDS-PAGE as described above. L4 is still notable for the single band above 55 kDa, and L33 for the high molecular weight signals. The high molecular weight signals in L4, which were not clearly visible in Fig. 3B, became evident (Fig. 3C). The band above 55 kDa was also evident in L33 (Fig. 3C). The signals of protein monomers present in L33 is in a similar range as that observed in L4 (Fig. 3C). In assessing line efficiency, we employed the PK1 line (Fig. 3C), a transgenic Physcomitrella line, which produces a synthetic complement regulator, MFHR1 (Top et al., 2019). MFHR1 also has the 8His-tag. The production yield of PK1 was 0.1 mg MFHR1/g moss fresh weight under non-optimised conditions (Top et al. 2019) and 0.82 mg/g fresh weight under optimised conditions (Ruiz-Molina et al. 2022). When considering both monomer and high molecular weight signals in L33, the signal strength of NTD-LhMaSp1-8Rep-CTD**-**8Htag is much higher than the PK1 signal strength (Fig. 3C). We extracted the NTD-LhMaSp1-8Rep-CTD**-**8Htag from double the amount of biomass, however, this potential yield of NTD-LhMaSp1-8Rep-CTD**-**8Htag protein may surpass the highest recorded amount of recombinant spider silk protein produced in plants to date, which stands at 0.08 mg/g tobacco leaf (Scheller et al. 2004).

To further confirm the successful production of NTD-LhMaSp1-8Rep-CTD**-**8Htag, we performed subsequent mass spectrometry (MS) analysis of the protein band in L4, and identified our target protein with a sequence coverage of 62 % (Fig. 4). The APsp was undetectable, indicating its correct cleavage. Detection of some amino acids at the beginning of the NTD and the last amino acids of the CTD confirm the production of the intact protein.

**Fig. 4.**
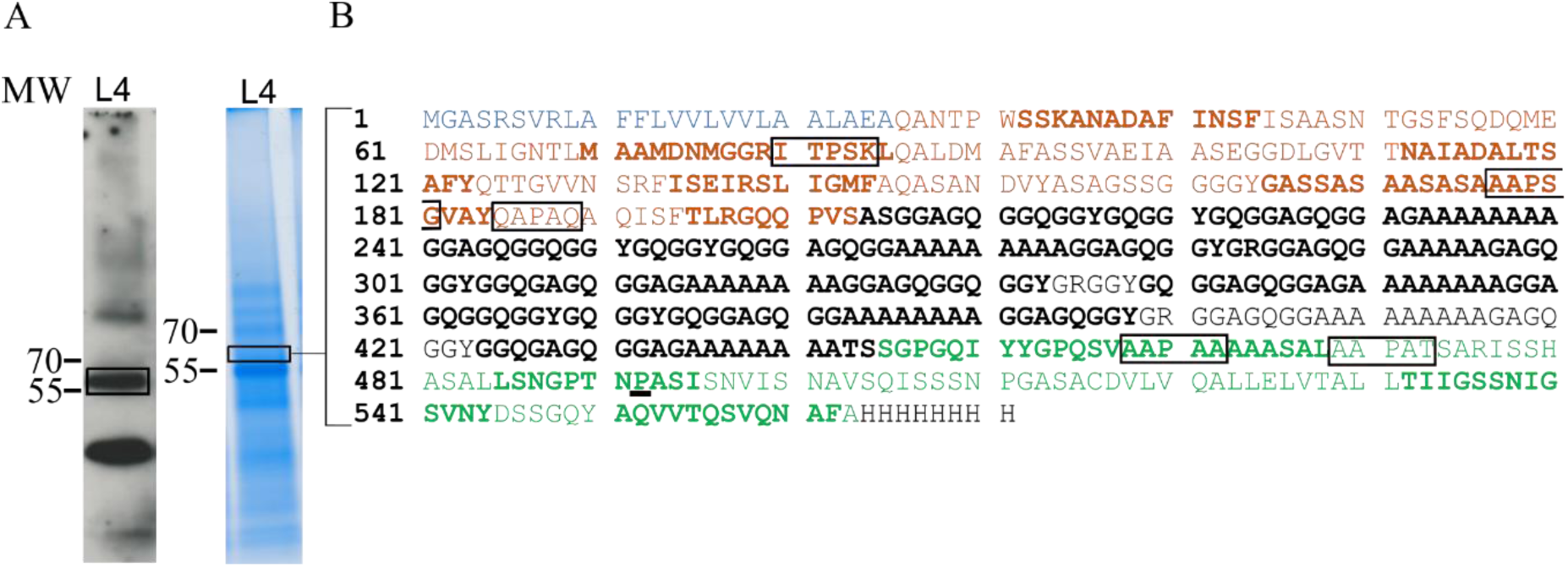
Mass spectrometric analysis of NTD-LhMaSp1-8Rep-CTD-8Htag. **A** Total protein of protonema material of L4 was recovered after acetone precipitation. The protein band at ∼56 kDa which is highlighted in the rectangle on the Coomassie-stained SDS-PAGE and corresponds to the band on the Western blot marked with a rectangle, was utilized for mass spectrometry. Anti His-tag antibody (18184, Thermo Fischer Scientific) was used for detection. **B** NTD-LhMaSp1-8Rep-CTD**-**8Htag peptides identified by mass spectrometry are shown in bold. Blue: Aspartic protease signal peptide (APsp). Orange: N-terminal domain. Black: repetitive poly-alanine and glycine blocks. Green: C-terminal domain. Potential sites for hydroxylation of proline are shown in rectangles. Identified hydroxyproline is underlined. Peptides for MS analysis were generated with chymotrypsin. Western blot under reducing conditions. MW: PageRuler Prestained Protein Ladder (Thermo Fisher Scientific).

One possible explanation for the high molecular weight signals is plant-typical *O*-glycosylation, which occurs on Physcomitrella proteins (Bohlender et al. 2022; Rempfer et al. 2024). To investigate this possibility, the JIM 16 antibody targeting arabinogalactans (Knox et al. 1991) was used. Arabinogalactans are a type of *O*-glycans, characterized by tree-like and multiple branched saccharide structures. Due to their size they can substantially elevate the molecular weight of proteins. Within the N-terminal domain of NTD-LhMaSp1-8Rep-CTD-H8tag, five potential sites for proline hydroxylation were identified (Fig. 4B). Hydroxylated prolines serve as the primary anchor for *O*-glycosylation such as arabinogalactans or extensin-like arabinosylation in plants (Gomord et al. 2010; Bohlender et al. 2022; Rempfer et al. 2024). Here, using MS analysis we did not find evidence for prolyl-hydroxylation or arabinosylation of three of these potential Hyp sites (IT**<u>P</u>**SK, AA**<u>P</u>**SG, AA**<u>P</u>**AA) whereas no peptide coverage for the other two sites (QA**<u>P</u>**AQ, AA**<u>P</u>**AT) was obtained (Fig. 4B). In contrast, hydroxylation of a C-terminal proline (TN**<u>P</u>**AS) residue was identified by MS, although at low confidence (Fig. 4B, and Supplemental Fig. S8). Moreover, no signals indicating the presence of arabinogalactans were detected in the investigated NTD-LhMaSp1-8Rep-CTD-H8tag protein containing samples (Supplementary Fig. S9), suggesting that *O*-glycosylation is absent despite present prolyl hydroxylation. This suggests that the presence of products with high to very high molecular weight is not due to the attachment of arabinogalactan sugar structures to the protein product but might be NTD-LhMaSp1-8Rep-CTD**-**8Htag multimers. It was reported before that higher order protein fibres can assemble *via* multimerization of medium-molecular weight proteins (Li et al. 2024).

Previous studies suggest that spider silk proteins can assemble under acidic pH conditions (Landreh et al. 2010; Hagn et al. 2011; Eisoldt et al. 2012). To test this possibility, we explored the influence of acidic and basic environments on the multimerization of NTD-LhMaSp1-8Rep-CTD-H8tag enriched *via* His-SpinTrap columns. Therefore, His-SpinTrap purified samples were adjusted to pH ranges of 2-3, 3-4, 5-6, 7-8, and 8-9, respectively. Subsequently, the samples underwent overnight incubation at 4°C and were analysed *via* Western blot and immunodetection. We found no noticeable differences in the very high molecular weight signals on the Western blot after incubation in acidic condition (pH 5-6) compared to basic conditions (Supplementary Fig. S10). Under extreme acidic pH conditions (pH 2-3, pH 3-4), the monomers were nearly absent. This might be because highly acidic pH leads to a decrease in monomer solubility, potentially resulting in protein precipitation in the sample. Moreover, degradation proteases are more active at acidic pH (Stael et al. 2019). A previous study reported the stability of a soluble 22 kDa *Araneus ventricosus* MaSp (monomers) at pH values from 2 to 12 for at least 1 h (Lee et al. 2012). Interestingly, extreme acidic pH did not lead to the precipitation of multimers, suggesting that these soluble multimers remain stable in extreme acidic environments. This stability implies that these multimers may have a robust structure that is resistant to pH changes. Our observations also reveal that within the basic pH ranges of 7-8 and 8-9, monomers and multimers are present without changes (Supplementary Fig. S10). It seems that variations in pH do not influence the conversion of monomers to multimers and *vice versa* under our conditions.

To assess the influence of other conditions on the multimerization of our protein, we tested varying pH levels within the growth medium (Supplementary Fig. S11), and different extraction buffers with different pH values (10, 7.4, 4.2) (Supplementary Fig. S12). None of these conditions prevented the formation of very high molecular weight soluble multimers. Furthermore, adding 75 mM DTT to the extraction buffer did not reduce the high molecular weight signals either (Supplementary Fig. S13). It was reported that some components in plant extracts, such as peroxidases, can induce cross-linking of coat proteins of the *Nudaurelia capensis* omega virus. (Castells-Graells and Lomonossoff 2021) They suggested cross-linking occurred during extraction and purification. However, their experiment of using extraction buffers with different pH, or adding 1 mM DTT did not make any difference to the result (Castells-Graells and Lomonossoff 2021).

The underlying cause of these notable soluble multimers remains unspecified within this study. We assume that the formation of these multimers can occur within the cell. The pH of the Golgi apparatus might have an impact on the formation of these multimers. The Golgi apparatus exhibits an acidic pH (6.8±0.2 to 6.3±0.3 from cis to trans Golgi in Arabidopsis) (Shen et al. 2013), the necessary pH for spider silk protein polymerisation, contrasting with the more basic pH of the ER (Shen et al. 2013). The soluble multimers observed in the NTD-LhMaSp1-8Rep-CTD**-**8Htag producing line could potentially stem from mature protein polymerisation due to the inherent property of proteins to polymerise in the Golgi apparatus, while the protein monomers may represent those still residing within the ER. This polymerisation can be beneficial for protein rearrangements necessary for the efficient *in-vitro* maturation of silk protein. However, if the soluble multimers stem from immature aggregates, either within the cell or during the extraction process, this could be detrimental to the protein rearrangements, thereby interfering with the maturation process *in vitro*.

The pH is not the only factor essential for spider silk protein polymerisation. Recently, a total of 180 metabolite components have been found in spider silk of *Trichonephila clavata* (Hu et al. 2023). Notably, the presence of two major metabolites, choline and DL-malic acid, within spider silk components highlights their pivotal roles and essential constituents in protein polymerisation and silk formation (Hu et al. 2023). Both components are also present in plant cells (George et al. 1934; Stafford 1956; Hanson et al. 1985; Andresen et al. 1988). As Physcomitrella is rich in secondary metabolites (Erxleben et al. 2012; Munoz et al. 2024), it may provide components for polymerisation of spider silk proteins.

### Purification of moss-produced NTD-LhMaSp1-8Rep-CTD-8Htag

For further analysis we purified the recombinant protein. We isolated His-tagged NTD-LhMaSp1-8Rep-CTD**-**8Htag from 9-day-old plant material cultivated under 2% CO_2_ and purified it using Ni-NTA chromatography. The target protein started eluting in fraction 19 at 140 mM imidazole, and was found in higher concentration in fraction 21 at 500 mM imidazole. Validation through Coomassie staining and Western blotting confirmed the presence of the expected monomers between 55 and 70 kDa (Supplementary Fig. S14 A, B). Western blot also shows multimers (Supplementary Fig. S14 B). The lower molecular bands are potential cross reaction signals from WT. Further optimisation is required to improve the purification process and eliminate unspecific bands. Further experimentation and method development are required to establish a robust and validated ELISA protocol for the quantification of the protein of interest in subsequent studies.

### Moss producing NTD-LhMaSp1-8Rep-CTD-8Htag has WT phenotype and similar biomass accumulation

We investigated the impact of NTD-LhMaSp1-8Rep-CTD**-**8Htag production on moss growth and phenotype. It was reported that the expression of MaSp2 is deleterious to *E. coli* through a negative effect on cell growth (Connor et al. 2023). Bioreactor cultures of Physcomitrella WT and NTD-LhMaSp1-8Rep-CTD**-**8Htag producing line L33 were started in parallel at a density of 50 mg DW/L (Fig. 5A). The microscopic analysis of protonema development observed in both lines at day 5 exhibited no differences (Fig. 5B). Biomass gain was monitored over a time period of 17 days by regular dry weight measurements. Biomass density in both lines reached almost 3 g DW/L (Fig. 5C) at day 17. The growth index (GI=biomass_final_**-**biomass_initial_)/biomass_initial_) for WT and L33 are 58 and 55.2, respectively. The results show that the biomass density and morphology of the production line was not significantly altered by LhMaSp1 protein production. The LhMaSp1 production (both monomers and soluble multimers) was confirmed in the moss line L33 growing in the photobioreactor (Fig. 5D).

**Fig. 5.**
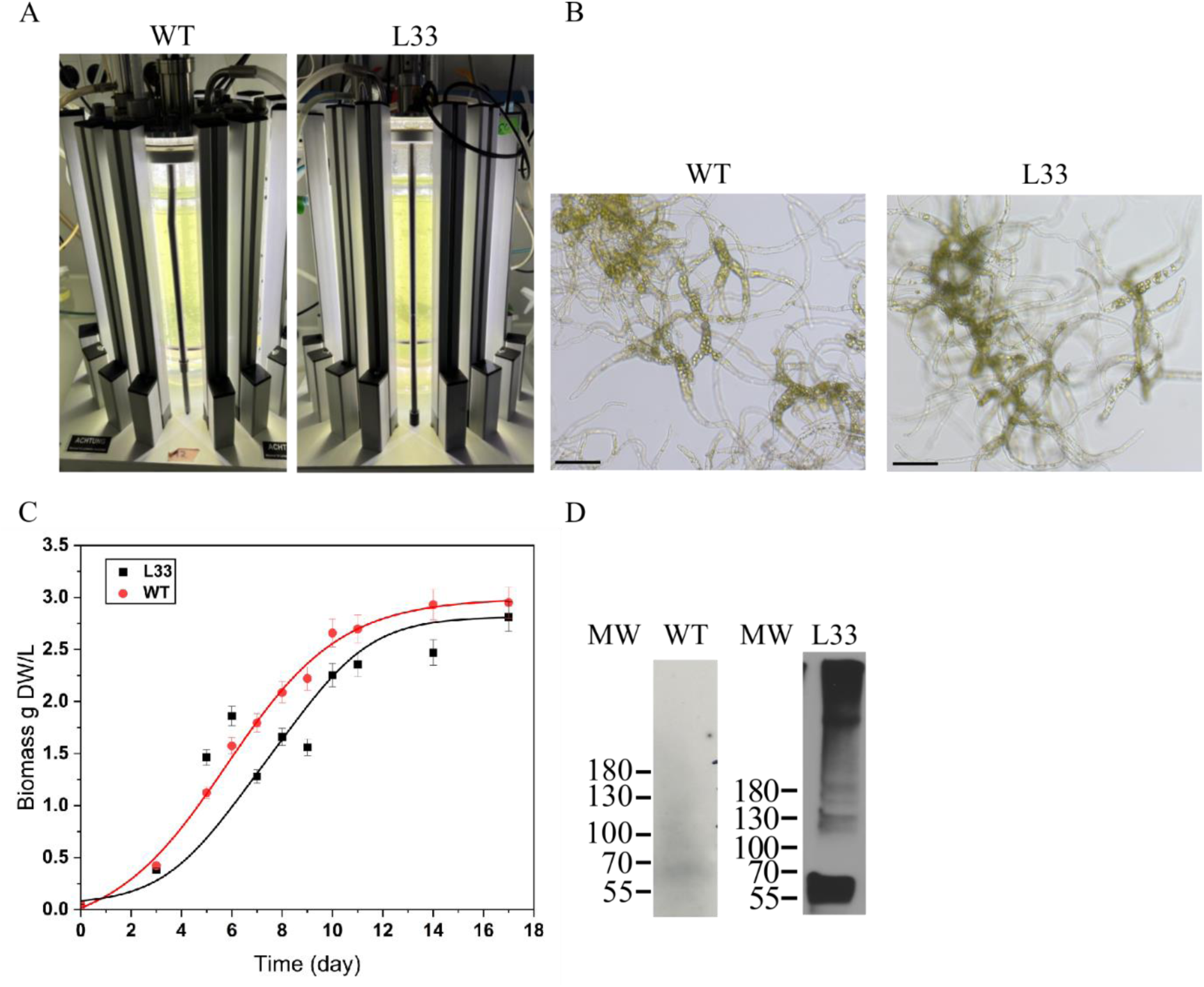
Cultivation of Physcomitrella WT and NTD-LhMaSp1-8Rep-CTD-8Htag producing line (L33) in photobioreactors. **A** Five-litre photobioreactor set-up of WT and NTD-LhMaSp1-8Rep-CTD**-**8Htag-producing line L33 was illuminated with LED lamps. After 2 days, the lights increased from 160 to 350 µmol/m^2^s. **B** Morphology of protonema tissue from WT and L33 at day 5 of cultivation. Bars = 150 µm. **C** Biomass accumulation of WT and L33 in the photobioreactor over a period of 17 days. Points represent mean values with error bars indicating standard deviation. WT (red points) and L33 (black points) showed exponential growth until day 8, after which the growth rate began to slow. Solid lines represent fitted curves for WT (red) and L33 (black). Statistical analysis showed no significant difference between the two lines (t-test, p > 0.05). Biomass density in both lines reached almost 3 g DW/L at day 17. **D** 100 mg biomass of both lines at day 8 was harvested for anti-His Western blot analysis. The result shows the production of NTD-LhMaSp1-8Rep-CTD**-**8Htag in both forms of monomers and soluble multimers. Anti His-tag antibody (18184, Thermo Fisher Scientific) was used. Western blot under reducing conditions. MW: PageRuler Prestained Protein Ladder (Thermo Fisher Scientific).

## Conclusion

In this study, we successfully achieved the stable production of recombinant NTD-LhMaSp1-8Rep-CTD**-**8Htag in Physcomitrella photobioreactors. Through genetic engineering and codon optimisation, we established a robust and consistent expression platform for NTD-LhMaSp1-8Rep-CTD**-**8Htag production in this moss. The estimated yield of Physcomitrella-produced LhMaSp1 is higher than 0.82 mg/1 g fresh weight, demonstrating the reliability and scalability of the platform. These results underscore the potential of Physcomitrella as an efficient host for the sustainable production of spider silk proteins, offering a promising avenue for the development of novel biomaterials and applications.

## Materials and methods

### Plant material

Physcomitrella wild type (new species name: *Physcomitrium patens* (Hedw.) Mitt.) was cultivated axenically in mineral KnopME medium 250 mg/L KH_2_PO_4_, 250 mg/L KCl, 250 mg/L MgSO_4_, 1 g/L Ca(NO_3_)_2_, 12.5 mg/L FeSO_4_, including microelements (50 μM H_3_BO_3_, 50 μM MnSO_4_ × H_2_O, 15 μM ZnSO_4_ × 7 H_2_O, 2.5 μM KJ, 0.5 μM Na_2_MoO_4_ × 2 H_2_O, 0.05 μM CuSO_4_ × 5 H_2_O, 0.05 μM CoCl_2_ × 6 H_2_O) according to Reski and Abel (1985), Egener et al. (2002) and Decker et al. (2015). LhMaSp1 producing lines were obtained by transformation of WT moss lines. Cultivation of Physcomitrella, protoplast isolation, polyethylene glycol-mediated transfection, regeneration and selection were performed according to Hohe et al. (2002), Hohe and Reski (2002) and Decker et al. (2015). Sterility of the culture media was monitored according to the protocol from Heck et al. (2021).

### Design of the vector NTD-LhMaSp1-12Rep-CTD-citrine for transient production

The CDSs of 12 repetitive poly-alanine blocks of *L. hesperus* silk protein core region (GenBank accession number EF595246), the full sequence of the N-terminal domain (GenBank accession number ABY67421.1) and the C-terminal domain (GenBank accession number AWK58707.1) were optimised (Supplementary Fig. S1) for Physcomitrella expression according to Top et al. (2021). This construct was synthesized in the pUC-GW-Amp plasmid by Genewiz (Leipzig, Germany). The CDS of NTD-LhMaSp1-12Rep-CTD was cloned into the expression vector containing PpActin5, the CDS of citrine, and the Nos terminator sequence *via Xho*Ⅰ and *Kpn*Ⅰ restriction sites. Gibson assembly (Gibson et al. 2009) was used to include the Physcomitrella aspartic protease 1 signal peptide (APsp; Schaaf et al. 2004). For this, the CDS of APsp was amplified from plasmid Act5_APsp_MFHR1 (Ruiz-Molina et al. 2022), together with the overhangs NTD and PpAct5 with the primers Fwd_overlapAct5_APsp and Rev_overlapNTD_APsp. The CDS of NTD was amplified from plasmid NTD-LhMaSp1-12Rep-CTD together with the overhangs APsp and 12-Rep region with the primers Fwd_overlapAPsp_NTD and Rev_overlap12Rep_NTD (Supplementary Table S1). 100 ng of each PCR product was added to the Gibson reaction including T5-Exonuclease, Phusion polymerase and Tag ligase (NEB, Frankfurt, Germany). The assembled vector was verified by sequencing. The plasmid (Supplementary Fig. S2A) is available *via* the International Moss Stock Center IMSC (https://www.moss-stock-center.org) under the accession number P2231 for the expression constructs APsp-NTD-LhMaSp1-12Rep-CTD-citrine. The protein sequence of NTD-LhMaSp1-12Rep-CTD is presented in Supplementary Fig. S2 B.

### Design of the vector NTD-LhMaSp1-8Rep-CTD-8Htag for stable production

The CDSs of 8 repetitive poly-alanine blocks of the *L. hesperus* silk protein core region (GenBank accession number EF595246), the full sequence of the N-terminal domain (GenBank accession number ABY67421.1) and the C-terminal domain (GenBank accession number AWK58707.1) were optimised (Supplementary Fig. S4) for Physcomitrella expression according to Top et al. (2021). The complete construct includes the Physcomitrella actin5 (P) promoter (Weise et al. 2006; Mueller et al. 2014; Niederau et al. 2024), 8His, a nos terminator, and a nptII (Neomycin Phosphotransferase II) antibiotic-resistance expression cassette. The intact construct was synthesized in the PUC-GW-amp plasmid by Genewiz (Leipzig, Germany). The APsp sequence was obtained from the plasmid APsp-NTD-LhMaSp1-12Rep-CTD *via* digestion with *Nhe*Ⅰ and *Eco*RV. The digested product was ligated to the NTD-LhMaSp1-8Rep-CTD**-**8Htag digested with the same enzymes. Assembled vectors were verified by sequencing. The plasmid (Supplementary Fig. S5A) is available *via* the International Moss Stock Center IMSC (https://www.moss-stock-center.org) under the accession number P2230 for the expression constructs APsp-NTD-LhMaSp1-8Rep-CTD-8Htag. The protein sequence of NTD-LhMaSp1-8Rep-CTD**-**8Htag is presented in Supplementary Fig. S5B.

### Confocal microscopy

To monitor the expression and localisation of NTD-LhMaSp1-12Rep-CTD-citrine proteins, confocal imaging was performed on transiently transfected protoplasts using a Zeiss LSM880 laser scanning confocal microscope (ZEISS, Jena, Germany). For all imaging experiments, an LD LCI Plan-Apochromat 40x/1.2 Imm AutoCorr DIC M27 water objective (ZEISS) was used with a zoom factor of 4. Citrine, chlorophyll, and mCerulean were excited with laser beams at 514 nm (Argon), 561 nm (DPSS), and 458 nm (Argon), respectively. The detection ranges were specified as 517-552 nm for citrine, 599-700 nm for chlorophyll, and 472-508 nm for mCerulean, respectively. The images were recorded in Zeiss ZEN black software and acquired as Z-stacks. For visual representation and analysis, the ZEN blue edition software was used. To produce three-dimensional reconstruction images, the maximum intensity projection option was used. To improve resolution, deconvolution was performed on all images.

### Plant screening and protein detection

For production of stable lines, plants surviving the neomycin selection as described in Decker et al. (2015) were transferred to liquid medium. After propagation of moss lines every two weeks for two months, they were subjected to screening for protein production. For this purpose, suspension cultures of 7-day-old protonema were vacuum-filtrated and 50-80 mg FW material transferred to 2-mL tubes with one tungsten carbide (Qiagen, Hilden, Germany) and one glass bead (Roth, Karlsruhe, Germany), 3 mm diameter. Plant material was homogenised with a tissue lyser (MM 400, Retsch, Haan, Germany) for 1 min at 30 Hz. Extraction buffer (50 mM Tris-HCl pH 7.2, 2% Triton X-100, 1% plant protease inhibitor cocktail P9599 from Sigma Aldrich) was added to the homogenized material and subjected to sonication for 15 min using an ultrasound bath (Sonorex RK52, Bandelin, Berlin, Germany), followed by centrifugation. Proteins were precipitated from the supernatant by adding 6x acetone allowing overnight incubation. Then pellets were dissolved in 50 mM Tris-HCl pH 7.2, 2% SDS at 95°C for 15 min. Before running in the SDS PAGE, 75 mM DTT and 4 x Laemmli sample buffer (Bio-Rad, Feldkirchen, Germany) were added. 50 µg of total protein (NanoDrop 1000 spectrophotometers, Thermo Fischer Scientific) was loaded on the SDS gel. Protein production was then analysed using Western blotting. For this, a 4-15% gradient gel (Mini-Protein TGX Precast Gels Bio-Rad) was run at 120 V for 1:30 h and blotted to polyvinylidene fluoride (PVDF) membrane (Hybond P 0.45, Amersham, Cytiva, Marlborough, USA) in a Trans-Blot SD Semi-Dry Electrophoretic Cell (Bio-Rad) for 1.15 h with 1.5 mA/cm^2^. The membrane was blocked for 1 h with 4% ECL blocking agent (Cytiva) followed by three times washing (1 x 15 min and 2 x 5 min) with Tris-buffered saline (TBST, 0.1% Tween 20). Then the membrane was probed with monoclonal anti-His antibody (18184, Thermo Fischer Scientific) as primary antibody overnight in 1:2000 dilutions with TBST buffer with 2% blocking agent. After three times washing, the secondary antibody, a sheep anti-mouse antibody coupled to horseradish peroxidase (NA931V, Invitrogen, Thermo Fisher Scientific, Massachusetts, USA), was diluted to 1:25,000 in TBST buffer with 2% blocking agent and incubated for 1 h on the membrane. After 4 times washing (2 x 10 min and 2 x 5min) the blot was incubated with detection reagents (SuperSignal West Pico PLUS Chemiluminescent Substrate, Thermo Fisher Scientific) and exposed to autography film (Hyperfilm ECL, Cytiva) up to 20 min.

### Rapid screening *via* His SpinTrap

The His SpinTrap column (Cytiva) was utilized for small-scale screening and purification of NTD-LhMaSp1-8Rep-CTD**-**8Htag proteins *via* immobilized metal ion affinity chromatography (IMAC). Protonema from suspension culture, cultivated for 9 days in a 500 mL Erlenmeyer flask under 2% CO_2_ in a shaker incubator, was harvested and 200 mg of biomass was homogenized using a tissue lyser. The material was then resuspended with 1.2 mL binding buffer (75 mM HNa_2_PO_4_.2H_2_O_2_ (disodium phosphate), 0.5 M NaCl, 20 mM imidazole, 0.05% Tween-20, 10% glycerol, 1% plant protease inhibitor cocktail (P9599, Sigma-Aldrich) pH 7 followed by 15 min sonication bath. After centrifugation, the supernatant was applied to the column previously equilibrated with binding buffer. The column was washed with 3x binding buffer followed by eluting the protein with 1x elution buffer (100 mM HNa_2_PO_4_x2H_2_O_2_, 0.5 M NaCl, 500 mM imidazole, 10% glycerol, pH 7.4). The eluted pool was collected in a new tube.

### Protein purification

NTD-LhMaSp1-8Rep-CTD**-**8Htag was extracted from vacuum-filtered plant material. For this, 4 mL binding buffer were added per gram FW and the suspension was disrupted with an ULTRA-TURRAX (Ika, Staufen, Germany) at 10,000 rpm for 10 min in an ice bath followed by sonication with an ultrasonic tip (Q500, QSONICA, Newtown, USA), amplitude 55%, volume <50 mL (time on: 10 sec, time off: 40 sec) for 20 min in total (on and off) on ice. Two consecutive centrifugation steps (4500 x g for 3 min and 20,000 x g for 10 min at 4°C) were carried out and the supernatant was filtered through 1 µm, and subsequently 0.22 µm PES filters (Roth, Dautphetal, Germany).

For chromatographic purification, the filtrate was loaded onto a 1 mL HisTrap FF column, using the ÄKTA system (GE Healthcare, Uppsala, Sweden) at 1 mL/min. The column was washed with 30 column volumes (CV) of binding buffer. The protein was eluted using a stepwise gradient (3% elution buffer for 9 CV, 15% 6 CV, 25% 3 CV, 100% 5 CV) (100%: 100 mM HNa_2_PO_4_.2H_2_O_2_, 0.5 M NaCl, 500 mM imidazole, 10% glycerol, pH 7.4), and collected in 1.5 mL fractions (fractions 9 to 23). The protein of interest was screened by Western blot and Coomassie staining.

### Mass spectrometry measurement and data analysis

Sample preparation for MS analysis was done as described in Hoernstein et al. (2016). In brief, gel slices at expected molecular weight were excised using a scalpel, chopped to pieces and destained with 30% acetonitrile (ACN), 70% 100 mM ammonium bicarbonate. Destained samples were shrunken in 100% ACN and dried in a speedvac. 0.2 µg chymotrypsin (Promega, Madison, USA) were applied per sample and proteolytic digest was performed for 16 h at 25 °C. Peptides were purified using C18-STAGE-Tips as described in Hoernstein et al. (2018) and eluted from the tip in 80% ACN, 0.1% formic acid (FA).

Measurements were performed on an Orbitrap Elite instrument (Thermo Scientific) coupled to an UltiMate 3000 RSLCnano HPLC system (Thermo Fisher Scientific) as described in Njenga et al. (2024). In brief, peptides were loaded on a nanoEase™ M/Z Symmetry C18 precolumn (20 mm x 180 µm ID; Waters) at a flowrate of 10 µL and separated on a µPAC column array (50 cm length; PharmaFluidics) at a flow rate of 0.3 µL/min at 40°C using a binary solvent gradient from 7% to 65% solvent B over 30 min with 0.1% (v/v) formic acid (FA) as solvent A and a mix of 0.1% (v/v) FA, 50% (v/v) methanol (MeOH), and 30% (v/v) ACN as solvent B. Subsequently solvent B was increased to 80% over 5 min and kept for 3 min. The HPLC was coupled online to Nanospray Flex ion source with the PST-HV as interface using a NFU liquid transfer system (MS Wil) and a fused silica emitter (EM-20-360; MicrOmics Technologies LLC).

MS^1^ scans were acquired at a mass range of m/z 370-1700 and a resolution of 120,000 (at *m/z* 400) using a target value (AGC) of 1 x 10^6^ ions, and a maximum injection time (IT) of 200 ms. MS^2^ scans were acquired from the top 15 ions (charge *≥* 2) and fragmentation was performed *via* collision-induced dissociation (CID) in the linear trap at a normalized collision energy of 35%, an activation time of 10 ms with a q-value of 0.25, a resolution of 35,000, an AGC target of 50,000 ions, a maximum injection time of 150 ms, and a dynamic exclusion time of 45 s. Raw data from MS measurements were processed with Mascot Distiller (V2.8.3.0, Matrix Science) and database searches were performed with Mascot Daemon (V2.6.0, Matrix Science) against a reverse concatenated database containing all Physcomitrella V3.3 protein models (Lang et al. 2018). In parallel, a custom database containing the sequences of known contaminants (e.g. keratin) was included. A precursor mass tolerance of ±10 ppm and a fragment mass tolerance of ±0.6 Da was used. Carbamidomethylation (C, + 57.021464 Da), deamidation (N, + 0.984016 Da), pyro-Glu formation (N-term Q, −17.026549 Da) and oxidation (M; P, + 15.994915 Da) were specified as variable modifications. Search results were loaded in Scaffold5 software (V5.0.1, Proteome Software) and protein hits were accepted at a false discovery rate (FDR) of 1% at the protein level and 0.5% at the peptide level while having at least two identified peptides.

### Bioreactor culture and determination of biomass

The cultures of WT and NTD-LhMaSp1-8Rep-CTD**-**8Htag producing line were scaled up to 5 L stirred tank photobioreactors (Applikon, Schiedam, The Netherlands) with Knop ME medium. Aeration 0.3 vvm (2% CO_2_), agitation with pitched 3 blade impeller at 500 rpm was used. After 2 days, the light was increased from 160 to 350 µmol/m^2^s. The pH level was kept constant at pH 5.8 by titrating with either 0.5 M NaOH or 0.5 M HCl, and the temperature was maintained at 22°C throughout the duration of the experiment. For dry weight (DW) measurement, 10–50 ml of tissue suspension was filtered and dried for 2 h at 105°C. Dry weight measurements were conducted for 17 days for both lines.

### Statistical analysis

The graph and analyses were performed with OriginLab software for windows (OriginLab, Northampton, Massachusetts, USA). Statistical significance was evaluated by t-test (p>0.05).

## Supplementary Information

The online version contains supplementary material available at (Publisher, insert URL).

## Acknowledgements

We thank the Life Imaging Center (LIC) of the University of Freiburg for expertise and support for confocal microscopy and Anne Katrin Prowse for language editing.

## Author contributions

M.R. designed research, performed experiments, analysed data, and wrote the manuscript. L.L.B. helped with construct design. J.P. helped with bioreactor experiments and with writing the manuscript. S.N.W.H. analysed the MS data, and helped writing the manuscript. E.L.D. supervised research and helped writing the manuscript. R.R. supervised research, acquired funding, and helped writing the manuscript. All authors read and approved the final version of the manuscript.

## Data availability

All datasets generated for this study are included in the manuscript and/or the Supplemental Files. The plasmid (Supplementary Fig. S2A) is available *via* the International Moss Stock Center IMSC (https://www.moss-stock-center.org) under the accession number P2231 for the expression constructs APsp-NTD-LhMaSp1-12Rep-CTD. The plasmid (Supplementary Fig. S5A) is available *via* the IMSC under the accession number P2230 for the expression constructs APsp-NTD-LhMaSp1-8Rep-CTD-8Htag. Lines 4 and 33 are available *via* the International Moss Stock Center IMSC under the accession numbers 40972 and 40973, respectively.

## Supplementary Information

**Fig. S1.**
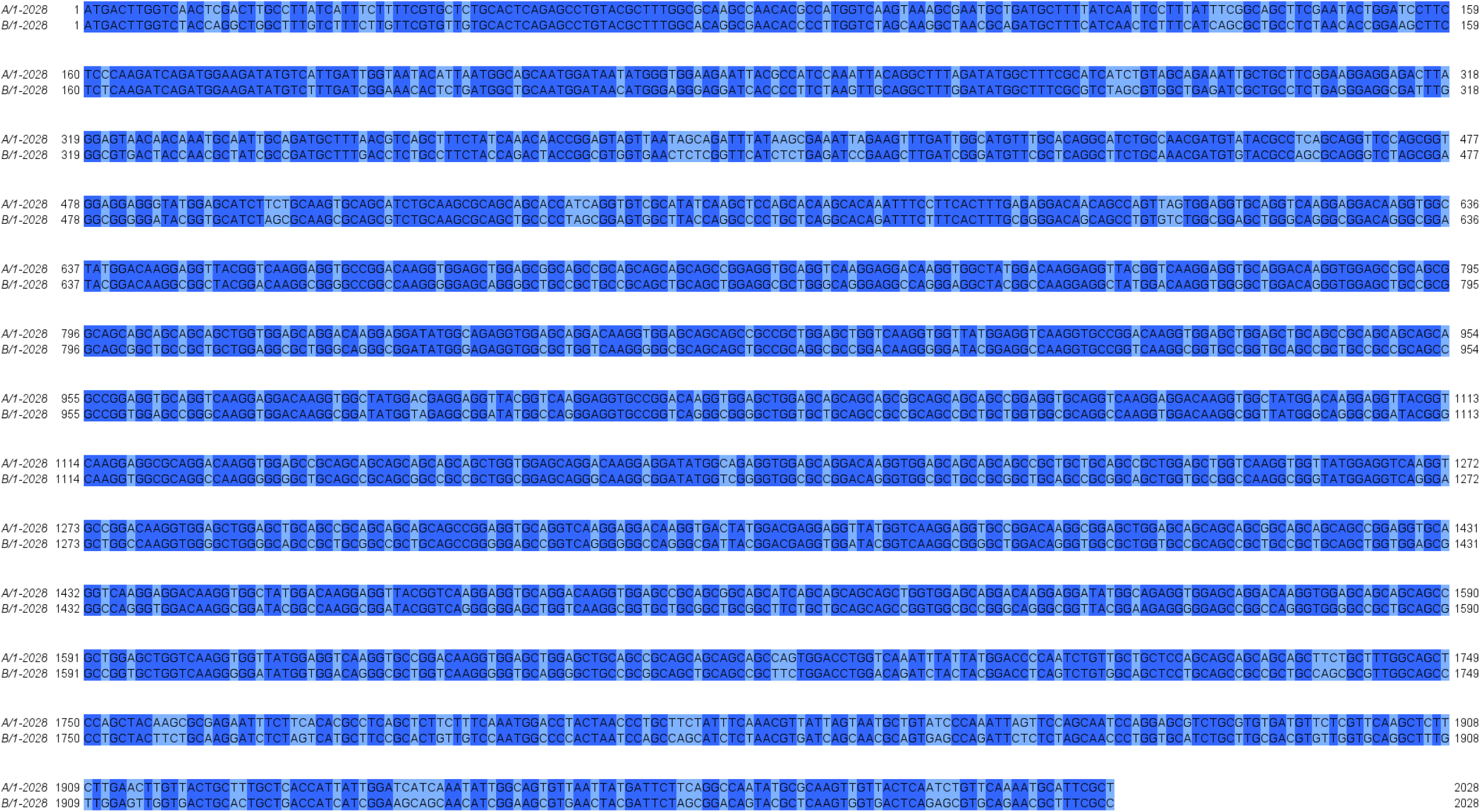
Pairwise sequence alignment of the original NTD-LhMaSp1-12Rep-CTD (A), and the codon optimized NTD-LhMaSp1-12Rep-CTD for codon usage in Physcomitrella (B). Alignment was performed with Jalview (Version 2.11.3.3, Waterhouse et al. 2009)

**Fig. S2.**
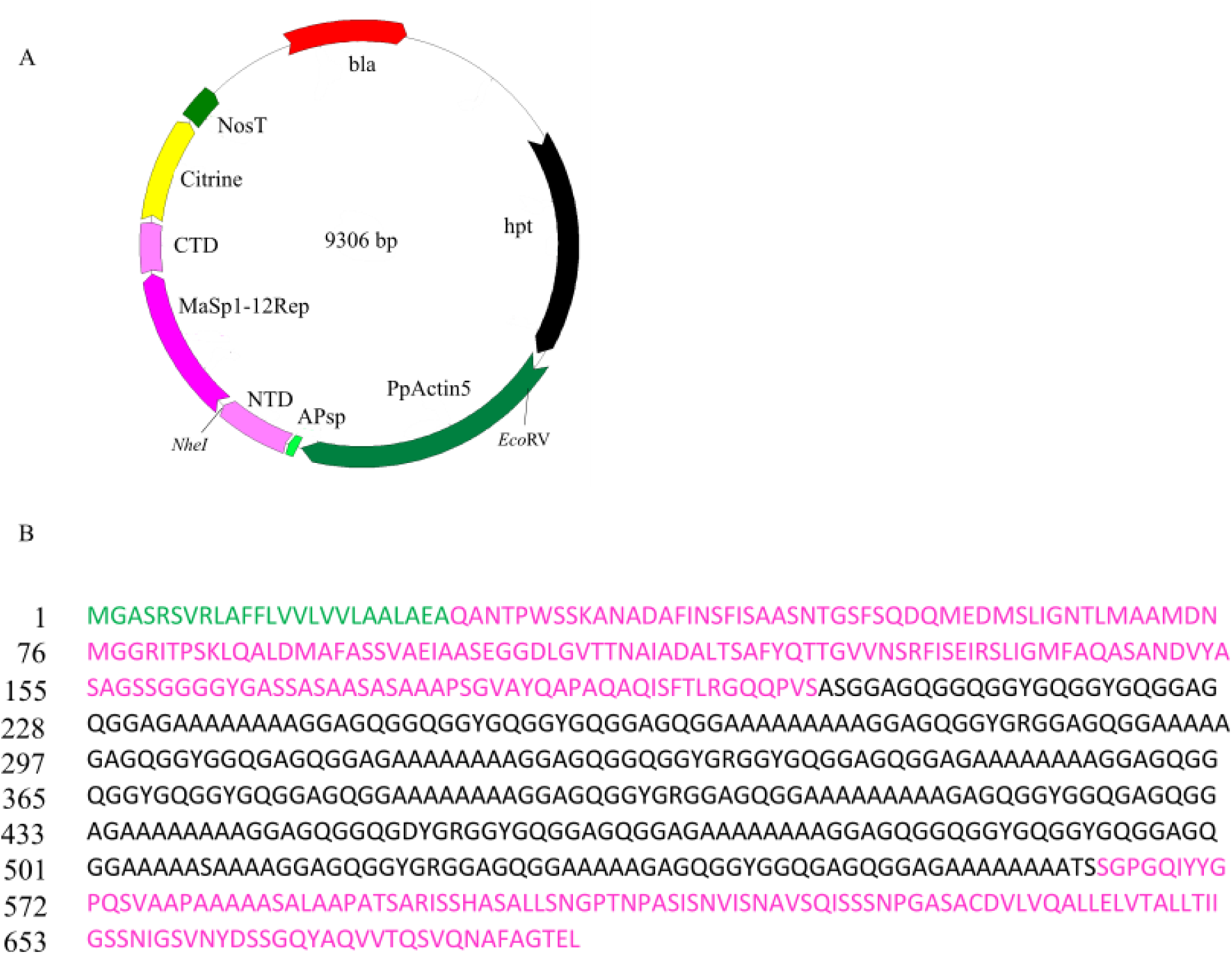
Designed construct for the transient expression of NTD-LhMaSp1-12Rep-CTD-citrine in Physcomitrella protoplasts. **A** Plasmid used for monitoring the localisation of MaSp1-12Rep into the secretory pathway of Physcomitrella. This plasmid includes the coding sequence of PpActin5 promoter, APsp, 12Rep, NTD and CTD, citrine Fluorescence protein, Nos terminator, and bla (beta-lactamase). **B** Protein sequence of NTD-LhMaSp1-12Rep-CTD. Green: APsp. Black: 12 repetitive Physcomitrella codon-optimized poly-alanine blocks of *L. hesperus* silk protein core region (12Rep). Purple: the full sequence of N and C-terminal domain (NTD and CTD).

**Fig. S3.**
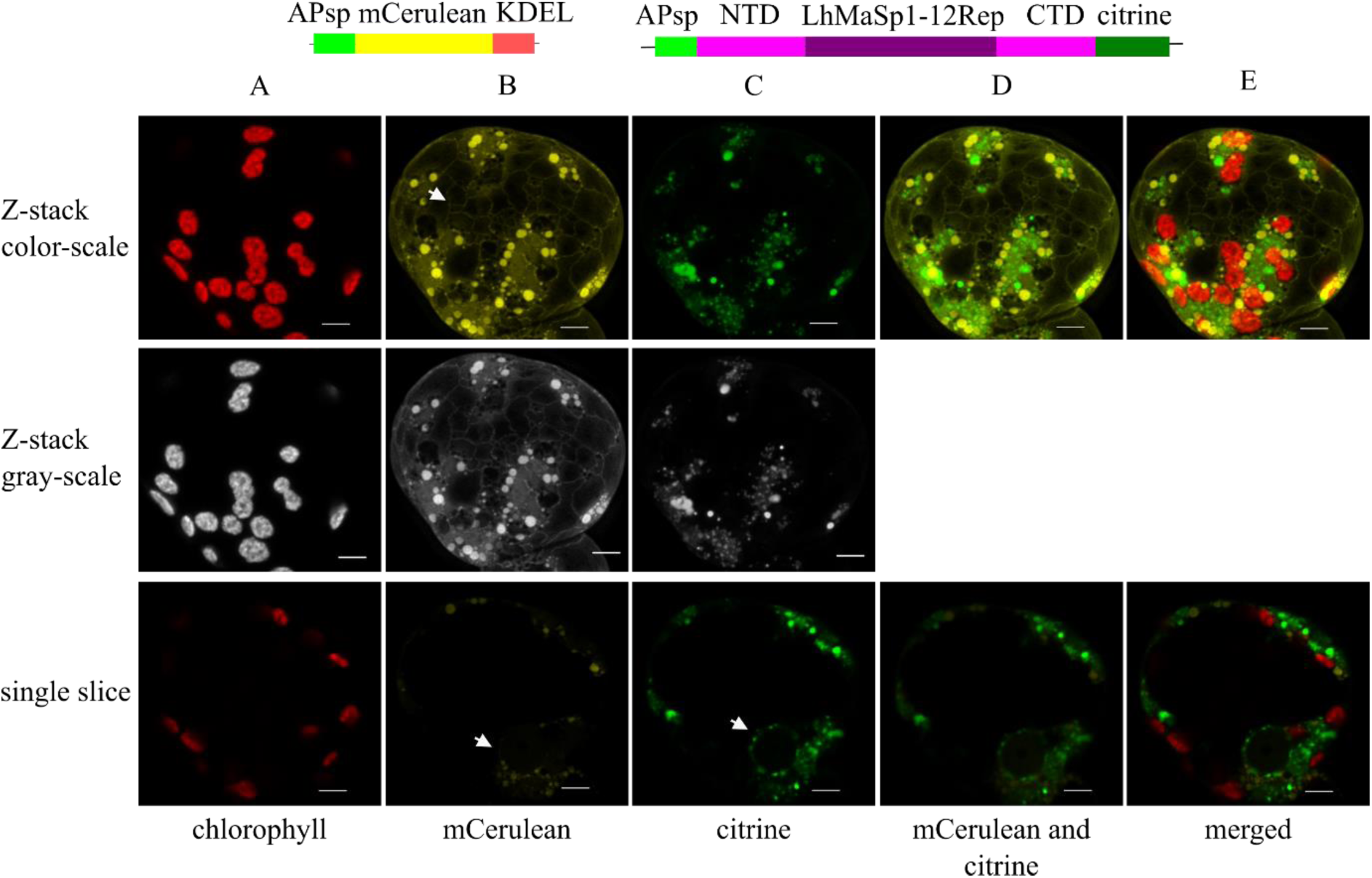
Confocal microscopy images showing the localisation of NTD-LhMaSp1-12Rep-CTD fused to citrine in Physcomitrella protoplasts. Schematic representation of the fusion proteins is at the top. The mCerulean-KDEL construct was used as positive control for ER localisation. **A** Chlorophyll autofluorescence. **B** The arrows show the fluorescence signal of ramified network of mCerulean-KDEL and its localisation to the nuclear envelope. **C** The arrow indicates the presence of NTD-LhMaSp1-12Rep-CTD protein fused to citrine in the nuclear envelope. The ramified network is not observed. **D** Citrine and mCerulean signals do not overlap, but cover almost the same area. **E** merged channels. The upper images display 3D-rendered Z-stack images, while the lower ones depict individual slices of a cell. Grey-scale channels are provided for better visibility of images. Bars = 5 µm.

**Fig. S4.**
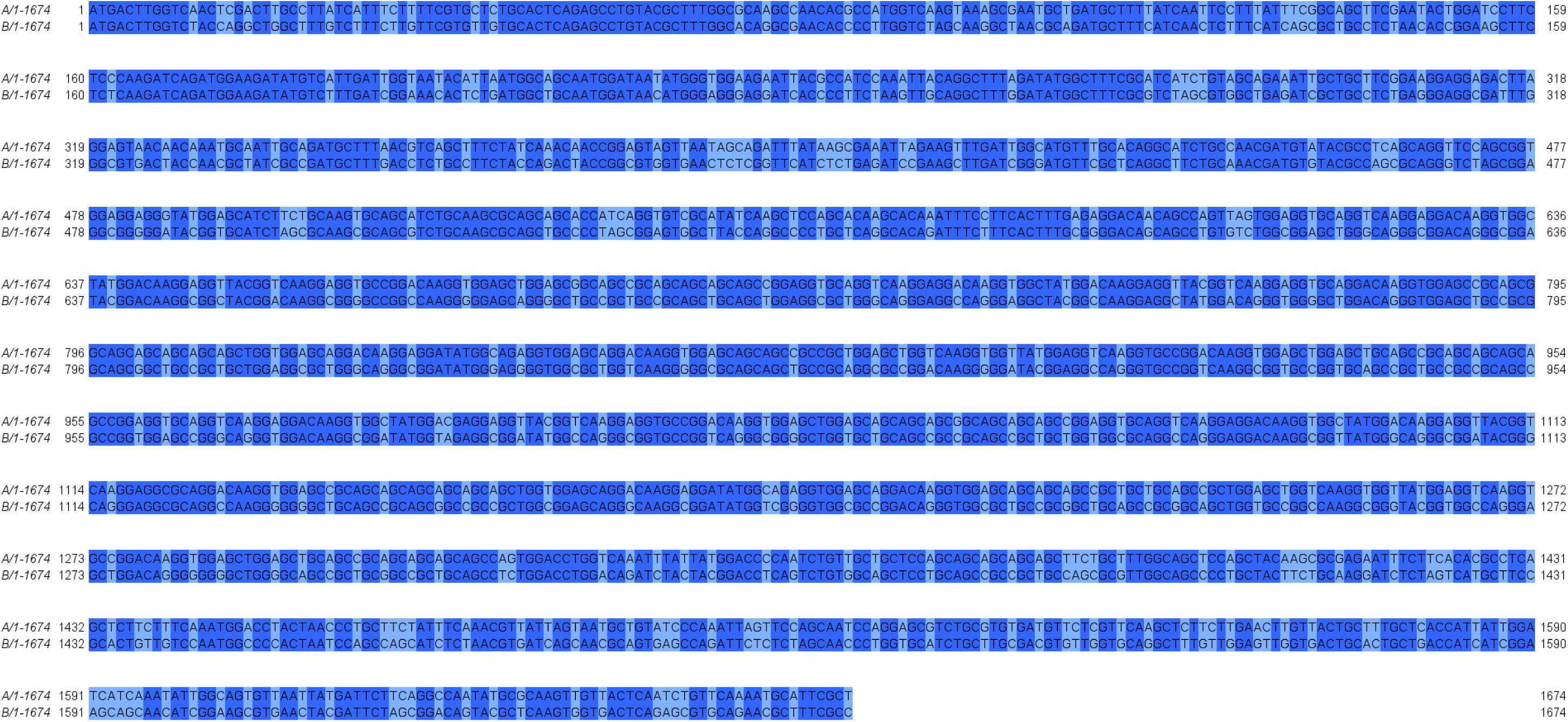
Pairwise sequence alignment of the original NTD-LhMaSp1-8Rep-CTD (A), and the codon optimized NTD-LhMaSp1-8Rep-CTD for codon usage in Physcomitrella (B). Alignment was performed with Jalview (Version 2.11.3.3, Waterhouse et al. 2009)

**Fig. S5.**
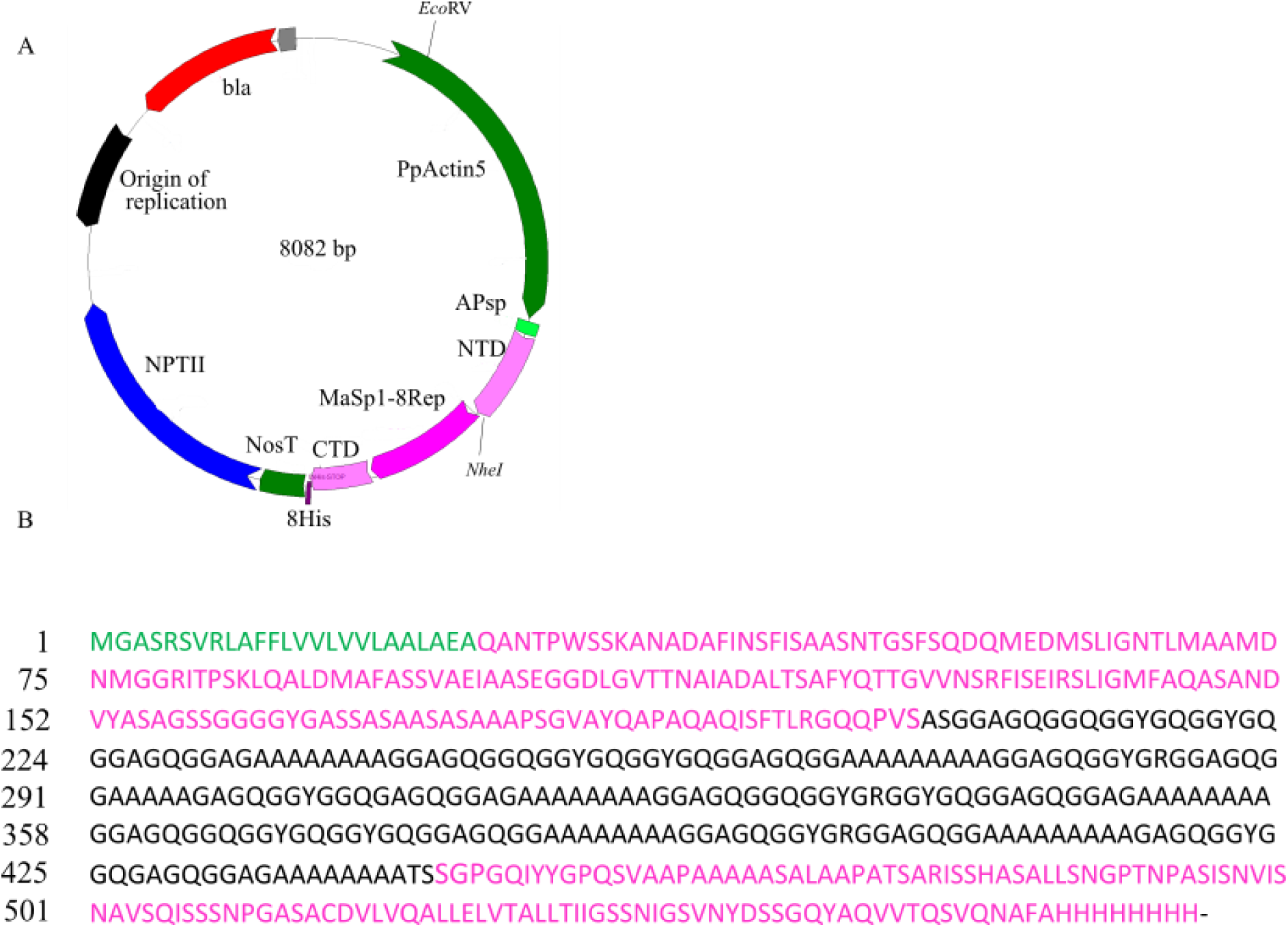
Designed construct for the stable production of recombinant NTD-LhMaSp1-8Rep-CTD-8Htag in Physcomitrella. **A** Plasmid for the stable NTD-MaSp1-8Rep-CTD**-**8Htag protein production. This plasmid includes the coding sequence of the Physcomitrella Actin5 promoter (PpActin5), APsp, 8Rep, NTD and CTD, 8His-tag, Nos Terminator, nptII (neomycin phosphotransferase) antibiotic-resistance expression cassette, and bla (beta-lactamase). **B** Protein sequence of NTD-LhMaSp1-8Rep-CTD**-**8Htag. Green: Aspartic protease signal peptide (APsp). Black: 8 repetitive Physcomitrella codon-optimized poly-alanine blocks of *L. hesperus* silk protein core region (8Rep). Purple: the full sequence of N and C-terminal domain (NTD and CTD).

**Fig. S6.**
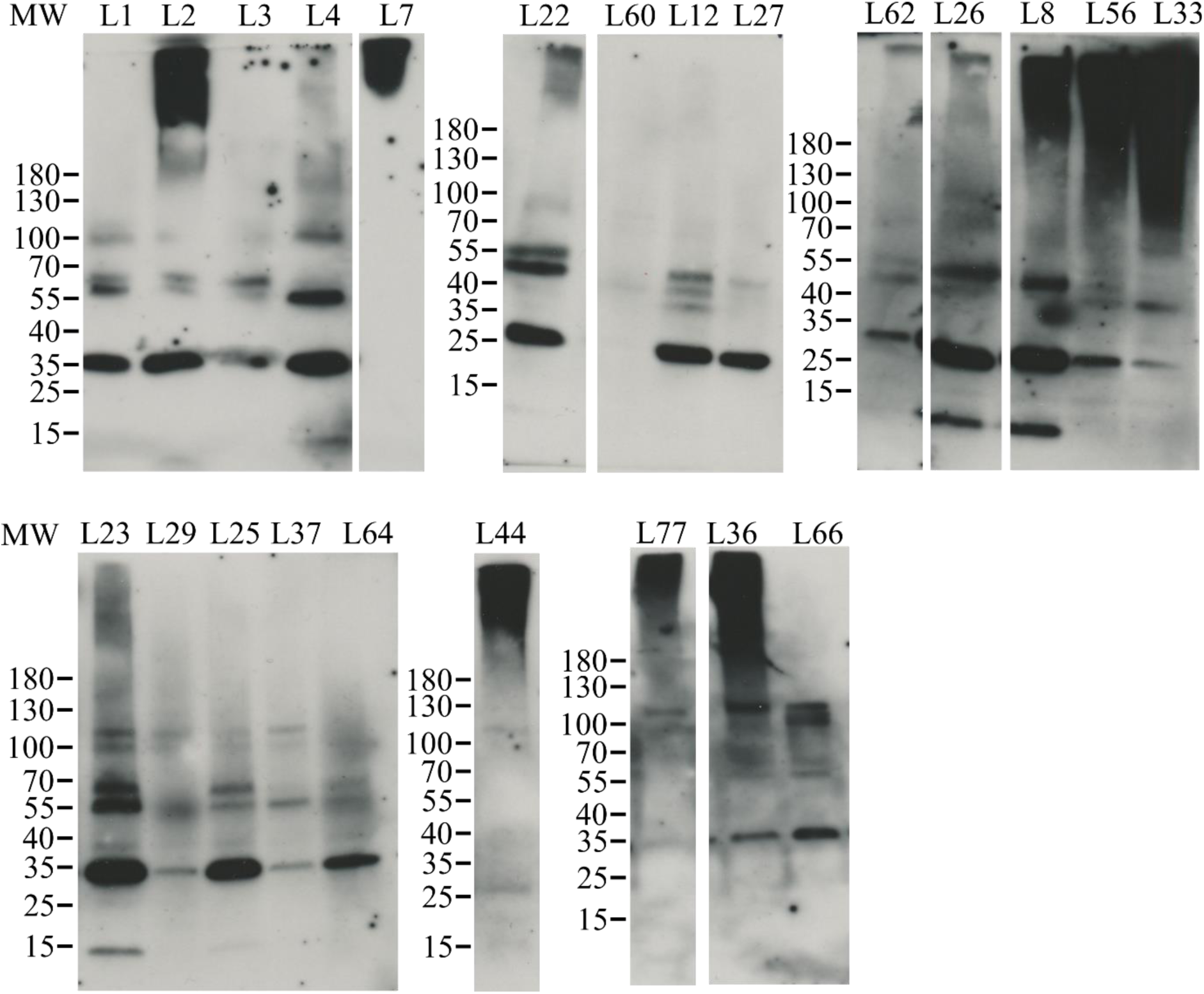
Western blot analysis of all 23 positive lines producing NTD-LhMaSp1-8Rep-CTD-8Htag. 50 µg of total protein was loaded for each line. Anti His-tag antibody (18184, Thermo Fischer Scientific) was used for detection. L: line. Western blot under reducing conditions. MW: PageRuler Prestained Protein Ladder (Thermo Fisher Scientific).

**Fig. S7.**
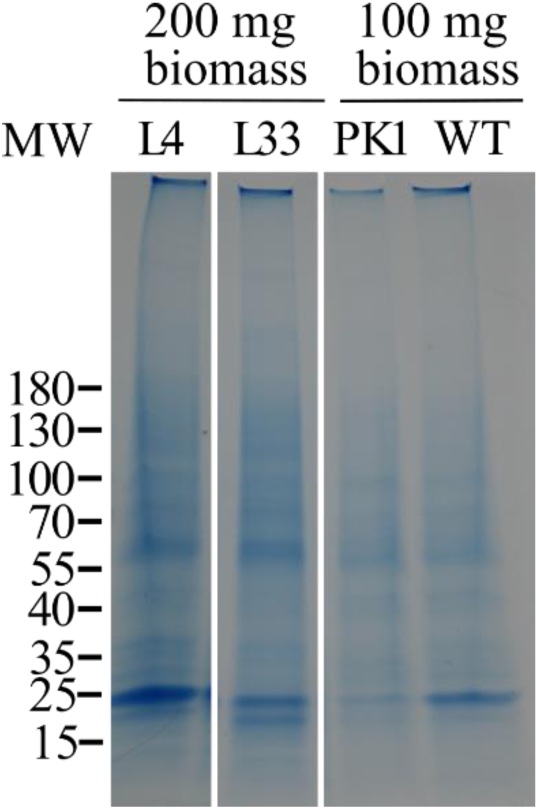
Coomassie staining as loading control for the total amount of protein loaded to each lane. Total soluble proteins extracted from 100 or 200 mg FW were subjected to His SpinTrap column, and 45 µl of the elution of L4, L33, PK1, and WT were loaded onto 4-15% gradient SDS-PAGE gels. PK1: transgenic moss line used as a positive control, as it efficiently produces a 58 kDa protein. L: line. MW: PageRuler Prestained Protein Ladder (Thermo Fisher Scientific).

**Fig. S8.**
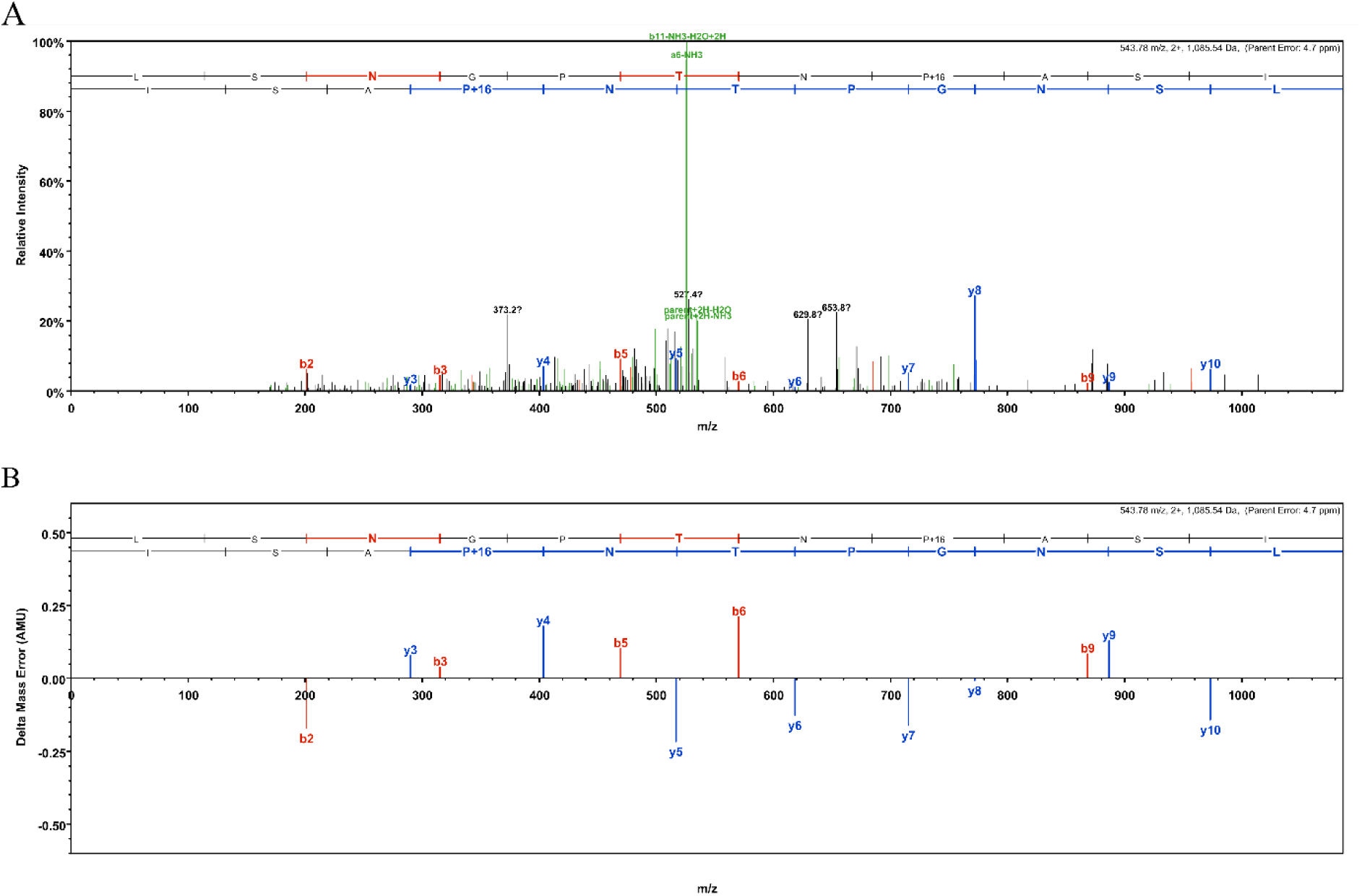
Collision-induced dissociation (CID) spectrum and fragment mass error distribution of the prolyl-hydroxylated peptide of MaSp1 recombinantly produced in Physcomitrella. **A** CID fragment mass spectrum of the peptide LSNGPTN**<u>P</u>**ASI. The mass shift of +16 indicates hydroxylation of the proline. **B** Fragment mass error distribution of b- and y-ions.

**Fig. S9.**
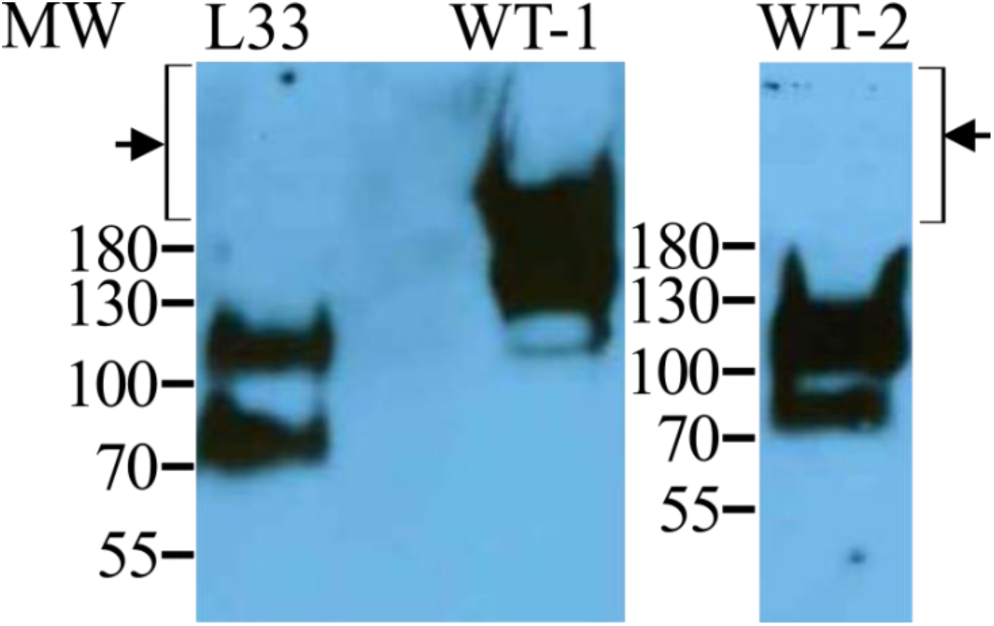
Western blot analysis of NTD-LhMaSp1-8Rep-CTD-8Htag, eluted through His SpinTrap column, using JIM 16 antibody against arabinogalactans. Total extracted proteins from wild-type (WT-1), and proteins of wild-type eluted from the His SpinTrap column (WT-2), were employed as positive controls, for arabinogalactan proteins. The arrow and bracket indicate the absence of arabinogalactans on NTD-LhMaSp1-8Rep-CTD**-**8Htag proteins, as well as negative control. Arabinogalactan proteins are present in the total extracted proteins of WT-1. 10 % SDS gels were used. Western blot under reducing conditions. MW: PageRuler Prestained Protein Ladder (Thermo Fisher Scientific).

**Fig. S10.**
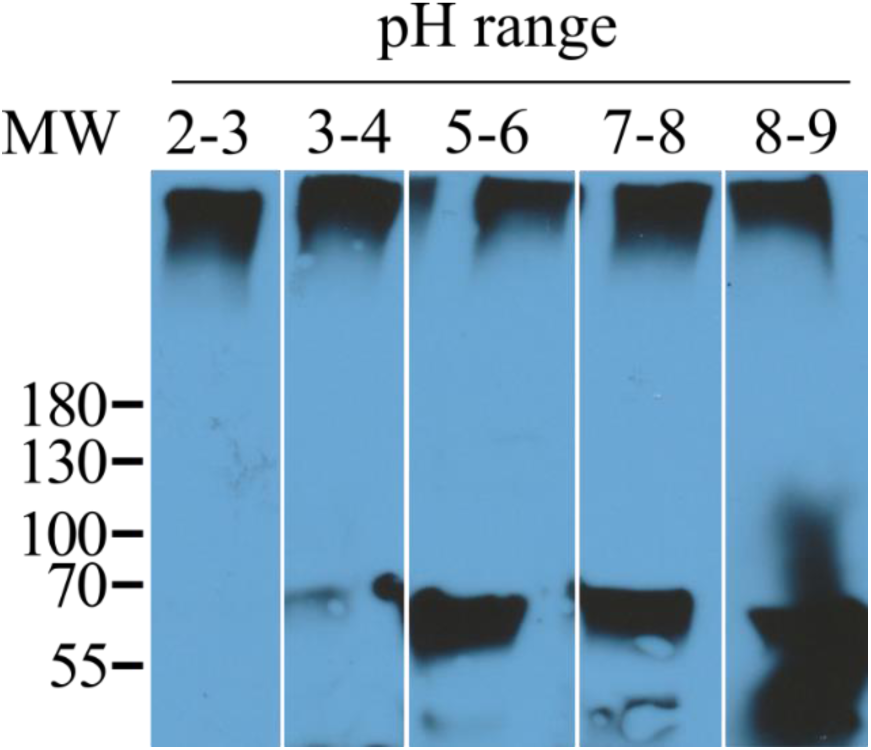
Effect of acidic and basic pH levels on the conversion of monomers to multimers and *vice versa*. Variations in pH do not appear to have a significant impact on the conversion of monomers to multimers and *vice versa*. A 10 % SDS gel was used. Western blot under reducing conditions. MW: PageRuler Prestained Protein Ladder (Thermo Fisher Scientific).

**Fig. S11.**
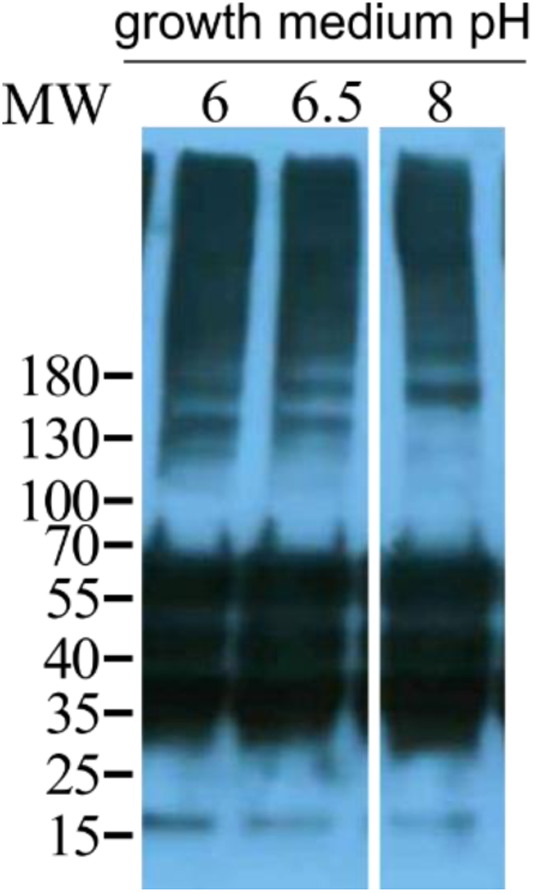
Effect of different growth medium pH on the high molecular weight signals. NTD-LhMaSp1-8Rep-CTD**-**8Htag extracted from L33 cultivated in acidic and basic medium with pH 6, 6.5 (buffered with MES), and 8 (adjusted with HEPES buffer), respectively. Western blot analysis showing that variations in the pH of the growth medium does not have any influence on the high molecular weight protein formation. Proteins at 40kDa, 35kDa, and 15kDa are unspecific signals also present in WT samples. MW: PageRuler Prestained Protein Ladder (Thermo Fisher Scientific). A 4-15% gradient SDS gel was used. Western blot under reducing conditions.

**Fig. S12.**
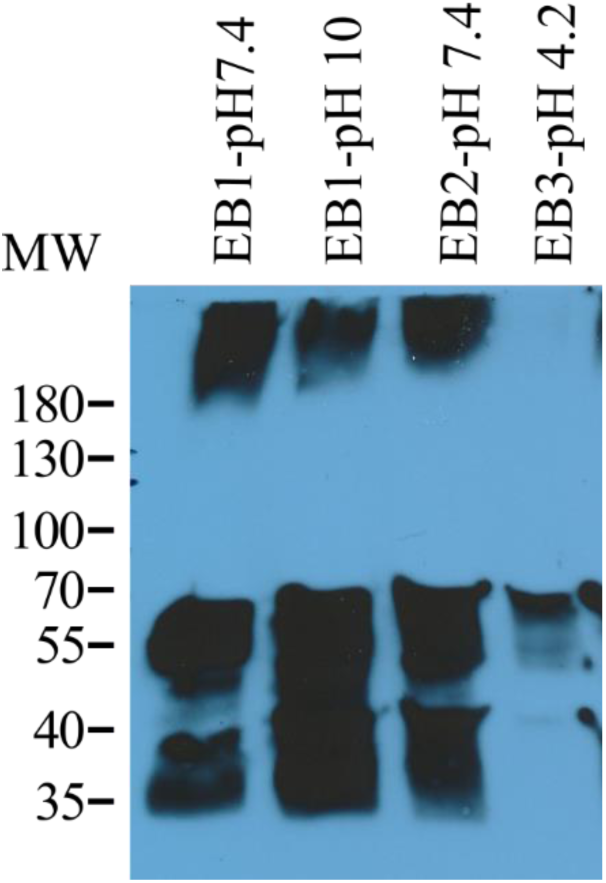
Effect of different extraction buffers (EB) and pH values on the high molecular weight signal. The high molecular weight signal was present for all extraction conditions except for pH 4.2. At pH 4.2, adequate extraction of total protein was not achieved. EB: extraction buffer. EB1: 75 mM HNa_2_PO_4_x2H_2_O_2_ (disodium phosphate), 0.5 M NaCl, 20 mM imidazole, 0.05% Tween-20, and 10% glycerol. EB2: 60 mM Tris, 0.5 M NaCl, 20 mM imidazole, and 10% glycerol. EB3: 60 mM MES, 0.5 M NaCl, 20 mM imidazole, and 10% glycerol. A 10% SDS gel was used. Proteins at 40kDa, and 35kDa are unspecific signals also present in WT samples. Western blot under reducing conditions. MW: PageRuler Prestained Protein Ladder (Thermo Fisher Scientific).

**Fig. S13.**
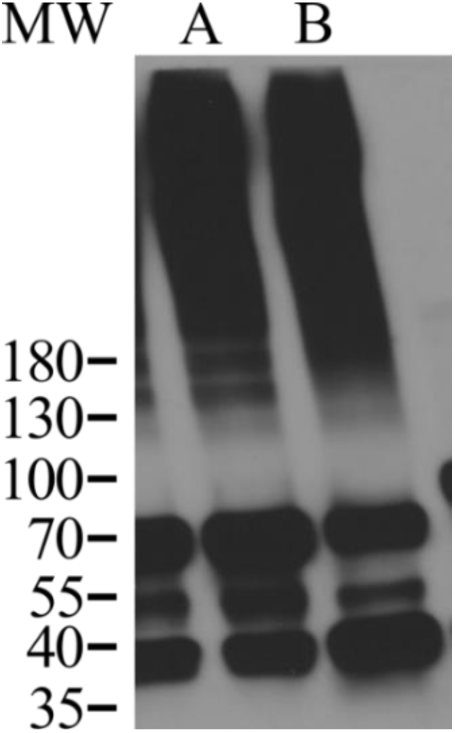
Influence of DTT in extraction buffer on the high molecular weight signals of NTD-LhMaSp1-8Rep-CTD-8Htag. **A** No DTT in extraction buffer. **B** Adding 75 mM DTT to the extraction buffer did not make a difference on signals from soluble multimer. Western blot under reducing conditions. MW: PageRuler Prestained Protein Ladder (Thermo Fisher Scientific).

**Fig. S14.**
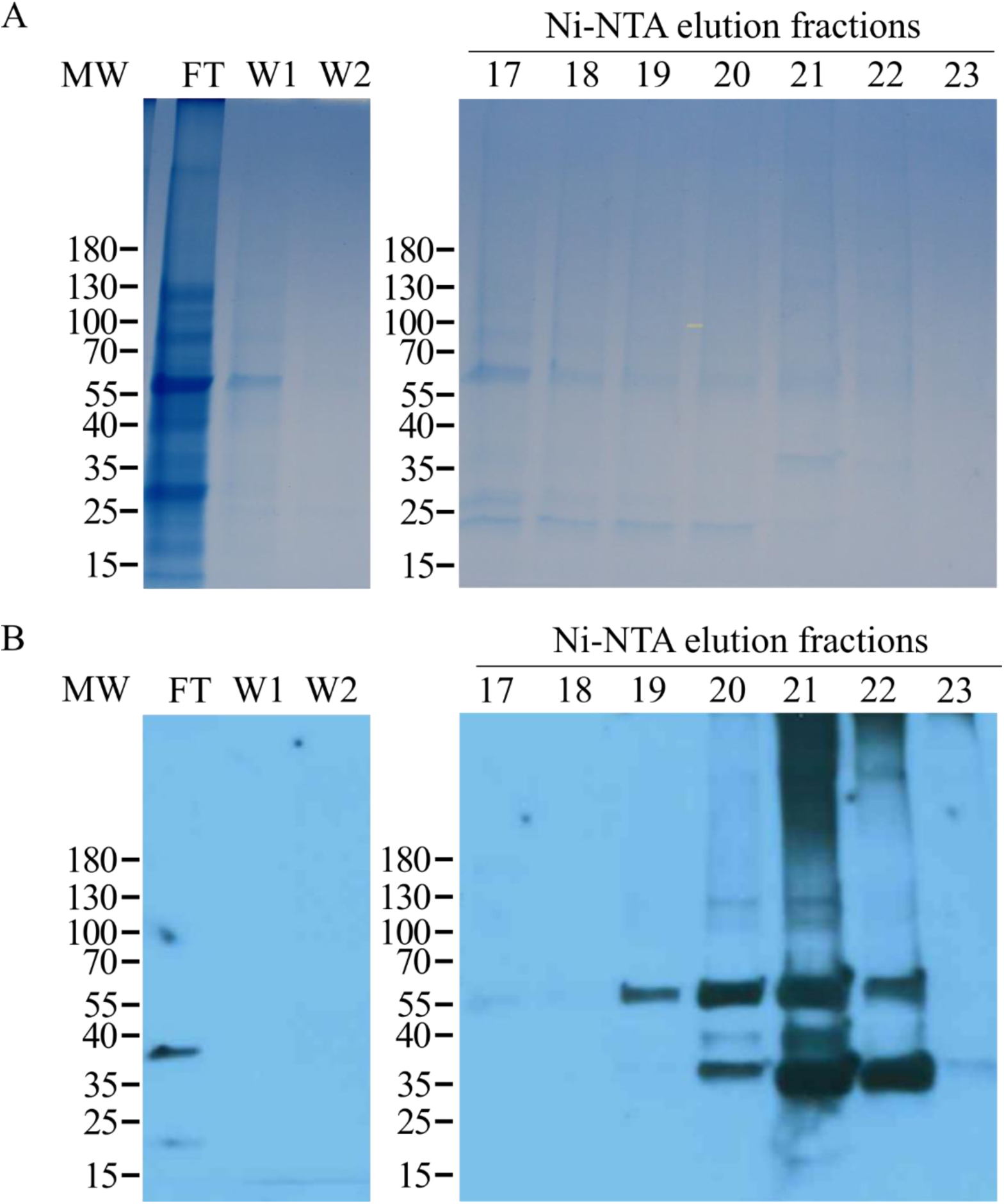
Purification of moss-produced NTD-LhMaSp1-8Rep-CTD-8Htag *via* Äkta Ni-NTA chromatography. **A** Coomassie staining of NTD-LhMaSp1-8Rep-CTD**-**8Htag elution fractions 17-23, Flow-Through (FT), wash1 (W1), and wash2 (W2). **B** Western blot of the same FT, W1, W2, and elution fractions detected with anti-His-tag antibodies. Western blot under reducing conditions. MW: PageRuler Prestained Protein Ladder (Thermo Fisher Scientific). A 4-15% gradient SDS gels were used.

**Table S1.**
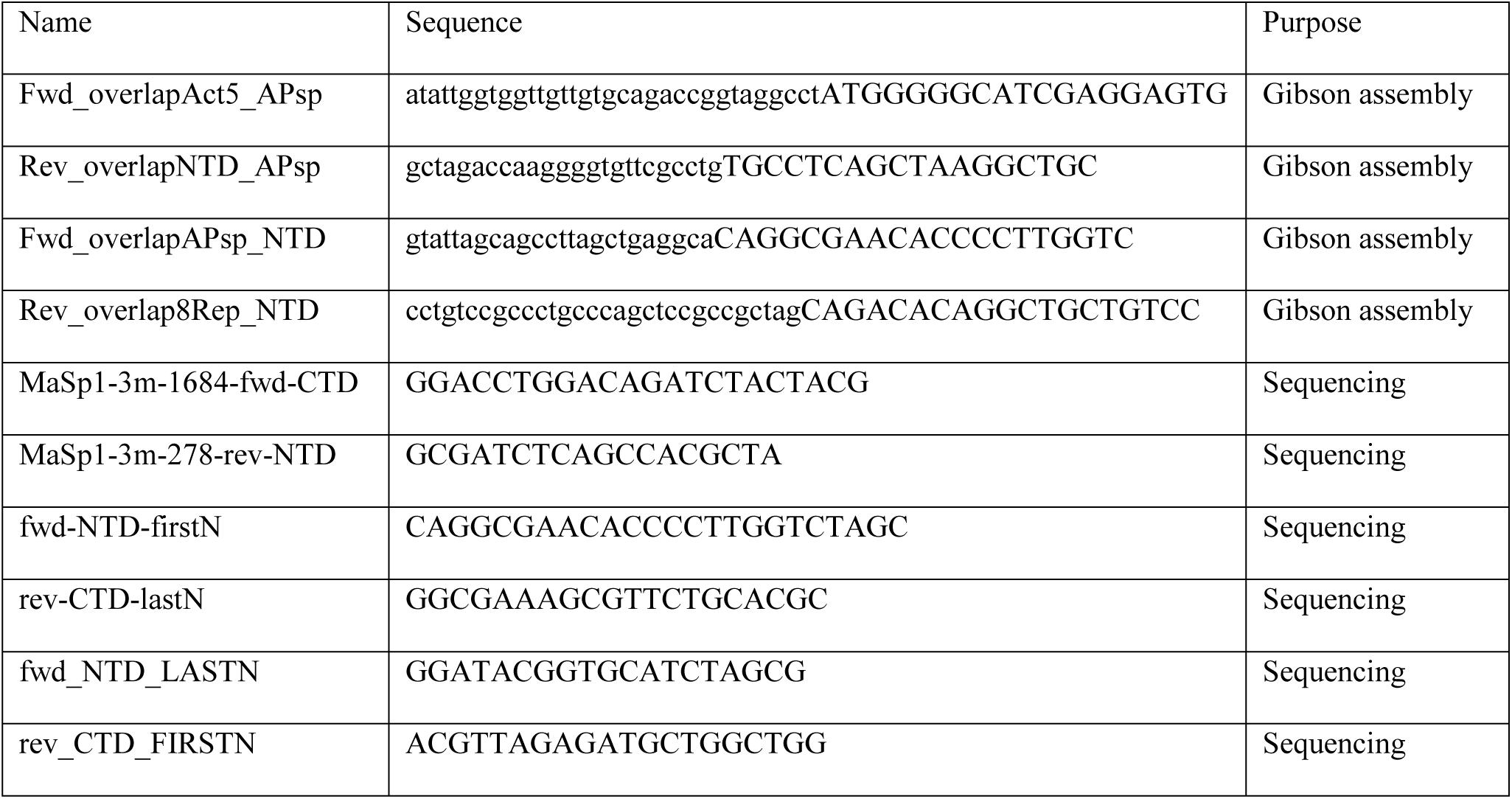
Oligonucleotides used for Gibson assembly and sequencing in this work. Overhangs are shown in lowercase.

